# Discovery and characterization of UCB-1A: the first PET radioligand for imaging synaptic vesicle glycoprotein 2C

**DOI:** 10.64898/2026.03.06.710176

**Authors:** Sangram Nag, Vasco C. Sousa, Anton Forsberg Morén, Miklós Tóth, Yasir Khani Meynaq, Elena Pedergnana, Rongfeng Zou, Anne Valade, Celine Vermeiren, Philippe Motte, Joël. Mercier, Xiaoqun Zhang, Per Svenningsson, Hans Ågren, Christer Halldin, Andrea Varrone

**Affiliations:** Department of Clinical Neuroscience, Centre for Psychiatry Research, Karolinska Institutet and Stockholm Health Care Services; Solna, Stockholm, SE-17164, Sweden; Department of Clinical Neuroscience, Division of Imaging Core Facilities, Autoradiography Core Facility, Karolinska Institutet; Solna, Stockholm, SE-17164, Sweden; Department of Physics and Astronomy, Division of X-ray Photon Science, Uppsala University; SE-75121 Uppsala, Sweden; UCB Biopharma, BE Research, Braine l’Alleud, Belgium; Department of Clinical Neuroscience, Neuro Section, Karolinska Institutet; Solna, Stockholm, SE-17164, Sweden

## Abstract

The synaptic vesicle glycoprotein 2C (SV2C) is a synaptic protein involved in the regulation of dopamine release. It is expressed in striatum, globus pallidus and substantia nigra, regions involved in the regulation of motor function. Genome-wide association studies, animal model and human brain tissue data indicate a strong link between SV2C and Parkinsońs disease, suggesting a potential role of SV2C as synaptic marker for Parkinsońs disease. We hypothesize that a positron emission tomography (PET) radioligand for SV2C can serve as imaging marker for Parkinsońs disease, enabling early diagnosis and assessment of disease progression. This study was therefore designed to develop a PET radioligand for imaging SV2C. UCB-1A was the lead candidate selected from a library of compounds developed by UCB BioPharma. A translational approach was used, including autoradiography and *in vitro* binding studies with [^3^H]UCB-1A, and in vivo PET studies with [^11^C]UCB-1A in non-human primates (NHPs). The *K*_D_ of [^3^H]UCB-1A for rat and human SV2C ranged between 6 and 15 nM, with >100-fold selectivity towards SV2A and SV2B. Specific binding of [^3^H]UCB-1A in rat and NHP brains was observed in substantia nigra, globus pallidus, striatum and brainstem nuclei, consistent with the expression of SV2C, and was decreased in the striatum of 6-hydroxydopamine-lesioned rats and in the putamen of Parkinson donors. UCB-1A was successfully radiolabelled with ^11^C and PET studies in NHPs demonstrated that [^11^C]UCB-1A displays suitable pharmacokinetic properties, a brain distribution consistent with the expression of SV2C and is selective for SV2C. [^11^C]UCB-1A is the first PET radioligand for *in vivo* imaging of SV2C and a potential synaptic marker for *in vivo* studies in Parkinsońs disease.

## INTRODUCTION

The synaptic vesicle glycoproteins belong to a family of proteins expressed in synaptic vesicles that are associated with other synaptic proteins involved in the regulation of synaptic release (*1*). There are three isoforms of SV2: SV2A which is highly abundant and uniformly distributed in the whole brain; SV2B which has lower density and uniform brain distribution; SV2C that has a specific expression in regions of the basal ganglia (*2, 3*). SV2C is expressed in dopaminergic neurons and neuropils in substantia nigra and striatum, in medium-sized spiny neurons in the globus pallidus, in cholinergic striatal neurons and in GABA-ergic terminals in substantia nigra pars reticulata (*4-6*). All are key regions of the basal ganglia system involved in motor control.

There is strong experimental evidence that suggests an important role of SV2C in Parkinson’s disease (*7*). First, polymorphisms of SV2C have been found to be associated with the protective role of nicotine and with L-dopa response (*7-10*). Second, genome-wide association studies have shown an association of SV2C with Parkinsońs disease in Asian and European populations (*11-13*). Third, studies in animal models have shown that SV2C: i) enhances dopamine retention in synaptic vesicles and modulates dopamine release (*14, 15*); ii) is decreased in the MPTP mouse model and is disrupted in the A53T Parkinsońs disease model (*15*); iii) has a protective effect on dopaminergic neurons in the MTPT mouse model (*14*); iv) co-immunoprecipitates with *a*-synuclein, suggesting an interaction between the two proteins (*15*). Finally, post-mortem studies in brain tissue from Parkinsońs disease donors have shown that the expression of SV2C is reduced in the substantia nigra (*16*) and that the protein is disrupted and colocalizes with ubiquitin in the striatum, indicating protein degradation (*15*).

In Parkinsońs disease, the accumulation of pathological aggregates of *a*-synuclein leads to synaptic dysfunction and down-regulation of synaptic proteins. These early changes, recapitulated under the definition of synaptopathy, are followed by subsequent degeneration of dopaminergic striatal terminals and neuronal loss (*17*). The availability of tools to study synaptic integrity and function is key for early detection of synaptopathy, paving the way for the development of Parkinsońs disease biomarkers for early diagnosis and assessment of disease progression. Molecular imaging of the dopaminergic system with positron emission tomography (PET) using radioligands for pre-synaptic targets is a well-established method to study the loss of striatal terminals in Parkinsońs disease (*18, 19*). More recently, with the development of [^11^C]UCB-J, a PET radioligand for imaging SV2A and synaptic density, PET studies have been conducted in different neurodegenerative diseases (*20, 21*). In Parkinsońs disease, a reduction of SV2A has been observed in the substantia nigra and locus coeruleus (*22, 23*). However, no longitudinal changes of SV2A have been observed in any brain regions, suggesting that SV2A PET imaging with [^11^C]UCB-J is not a suitable synaptic imaging marker to monitor disease progression (*22, 24*). We therefore hypothesize that, considering the enrichment of brain distribution of SV2C in the basal ganglia and the evidence of an interplay between SV2C and Parkinsońs disease, a PET radioligand for imaging SV2C would be an imaging marker for early detection of synaptopathy in Parkinsońs disease and a synaptic marker to monitor disease progression.

This study was designed to discover a PET radioligand for SV2C. A translational approach was used consisting of the following steps. First, from a series of compounds developed by UCB, we identified UCB-1A as potential candidate for further development as PET radioligand. Second, UCB-1A was radiolabelled with ^3^H and was characterized in vitro on recombinant paralogues of SV2, on tissue slices and homogenates from wild type rat brain, and on tissue slices from non-human primate (NHP) brain. Third, specific binding of [^3^H]UCB-1A was measured in brain slices from striatum and substantia nigra of 6-hydroxydopamine lesioned rat model and from post-mortem human brain samples, from donors with Parkinsońs disease. Finally, radiolabelling of UCB-1A with ^11^C was developed and the pharmacokinetic properties and in vivo selectivity of [^11^C]UCB-1A was evaluated with PET studies in NHPs.

## RESULTS

### Discovery of UCB-1A

From a library of 180 000 compounds available at UCB BioPharma, approximately 4000 compounds with non-racetam structure were preliminary selected. From these, two selective hits were identified (**Table S1**). From the first hit, two compounds (1a and 1b) displayed an in vitro pIC_50_ for human SV2C of 6.8 (1a) and 7.4 (1b). From the second hit, seven compounds were identified (**Table 1**) with pIC_50_ for hSV2C between 7 and 8.3 and a pIC_50_ for hSV2A between 5 and 7.2. Based on the affinity for hSV2C, the relative selectivity (hSV2C/hSV2A=1.15), logP<2 and logD<3, as well as the feasibility of labelling with ^3^H and ^11^C, UCB-1A was selected for further in vitro and in vivo evaluation.

**Table 1.**
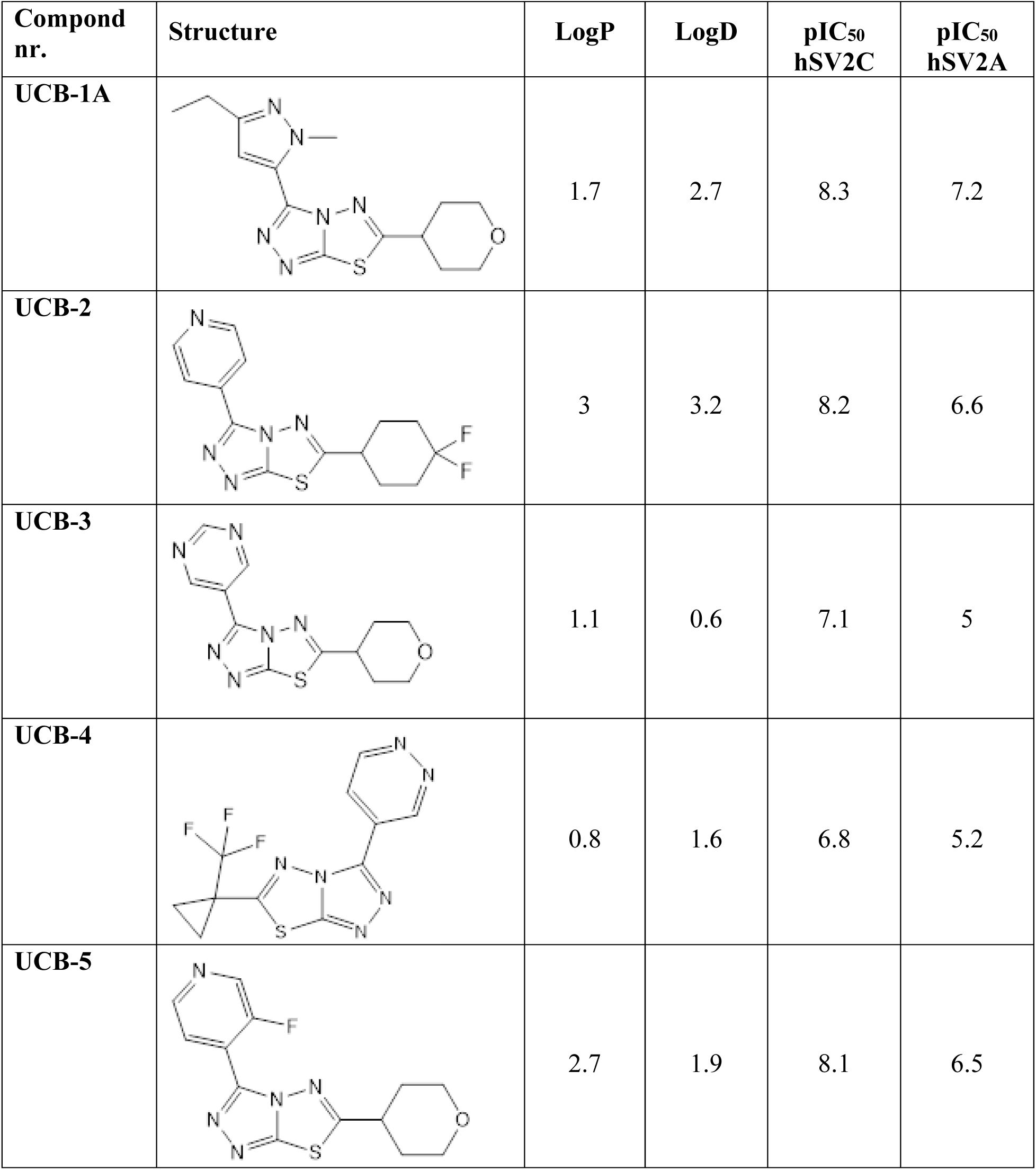
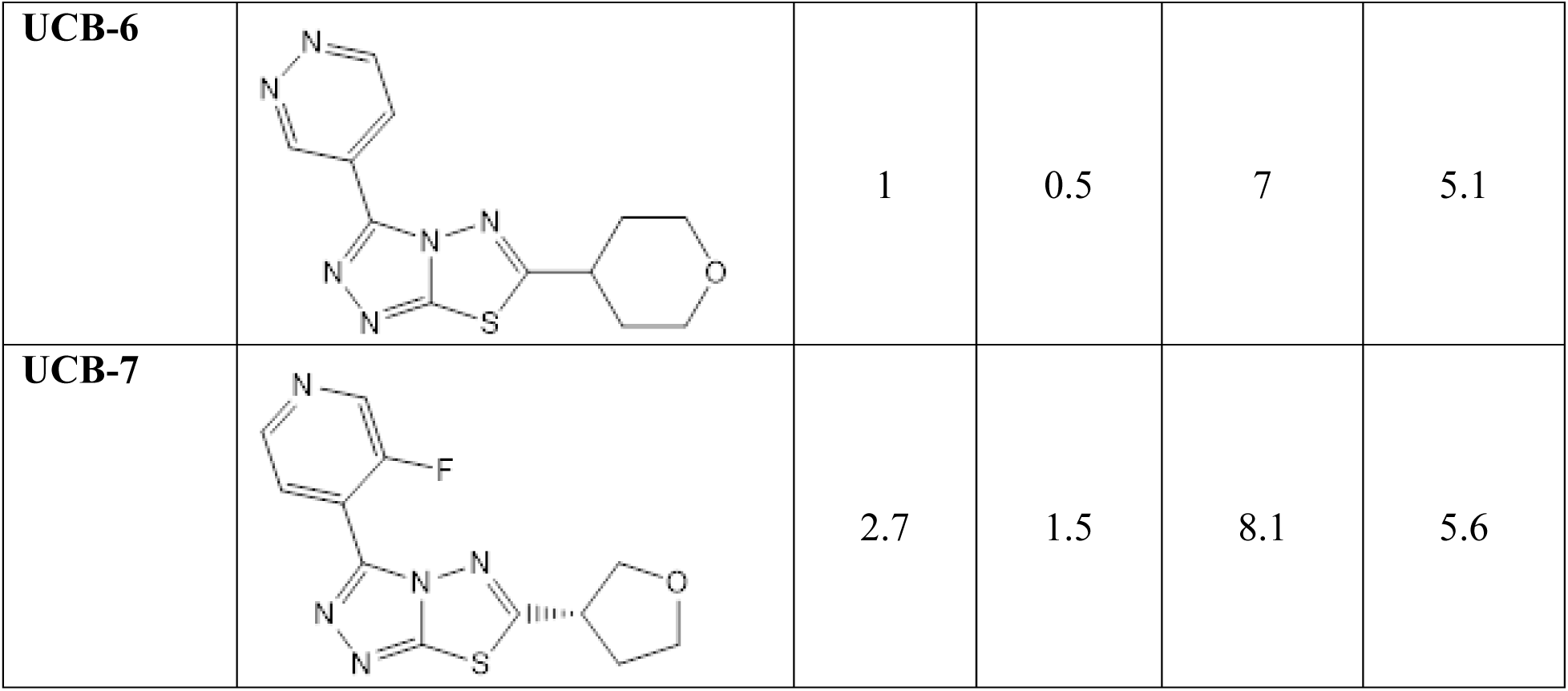
Physicochemical properties of UCB compounds.

| Compound nr. | Structure | LogP | LogD | pIC<sub>50</sub> hSV2C | pIC<sub>50</sub> hSV2A |
| --- | --- | --- | --- | --- | --- |
| UCB-1A |  | 1.7 | 2.7 | 8.3 | 7.2 |
| UCB-2 |  | 3 | 3.2 | 8.2 | 6.6 |
| UCB-3 |  | 1.1 | 0.6 | 7.1 | 5 |
| UCB-4 |  | 0.8 | 1.6 | 6.8 | 5.2 |
| UCB-5 |  | 2.7 | 1.9 | 8.1 | 6.5 |
| UCB-6 |  | 1 | 0.5 | 7 | 5.1 |
| UCB-7 |  | 2.7 | 1.5 | 8.1 | 5.6 |

### In silico modeling of UCB-1A binding to SV2 proteins

To characterize the binding mode of UCB-1A across SV2 paralogues, in silico modeling of the ligand bound to SV2A and SV2C was performed. A highly conserved pose within the binding pocket of all three paralogues appeared to be adopted by UCB-1A. The ligand appeared to be anchored by two principal interactions: (i) a π–π contact between its aromatic core and a nearby aromatic residue within the pocket, and (ii) a hydrogen bond between a protonated Asp side chain and a polar substituent of UCB-1A, with Asp acting as the hydrogen-bond donor. These recurring interactions define a common structural framework for ligand recognition within the SV2 family. Simulations conducted at 37°C revealed that the stability of UCB-1A binding to SV2A was lower than the one observed at 4°C, whereas binding to SV2C was comparatively stable at both temperatures (details of these simulations will be reported separately, manuscript in preparation).

### In vitro determination of K_D_ of [^3^H]UCB-1A on recombinant protein and rat brain homogenates

In vitro saturation binding studies were performed at UCB BioPharma on recombinant SV2 paralogues at 37°C using [^3^H]UCB-1A ([^3^H]UCB4297). The *K*_D_ of [^3^H]UCB-1A for human, non-human primate and mouse SV2C was 6 nM, 7.5 nM, and 15 nM. The *K*_D_ of [^3^H]UCB-1A for human SV2A and SV2B was ∼400 nM and ∼800 nM.

In a separate set of experiments conducted at UCB BioPharma, it was shown that the affinity of UCB-1A for SV2C was similar at 4°C and 37°C and confirmed the >100 fold selectivity of UCB-1A for SV2C vs. SV2A and SV2B (**Table S2**). *In vitro* saturation binding studies were performed at KI on rat brain homogenates from basal ganglia and brainstem (SV2C high-density regions) and cortex, hippocampus and cerebellum (SV2C low-density regions) (**Fig. 1**).

**Figure 1.**
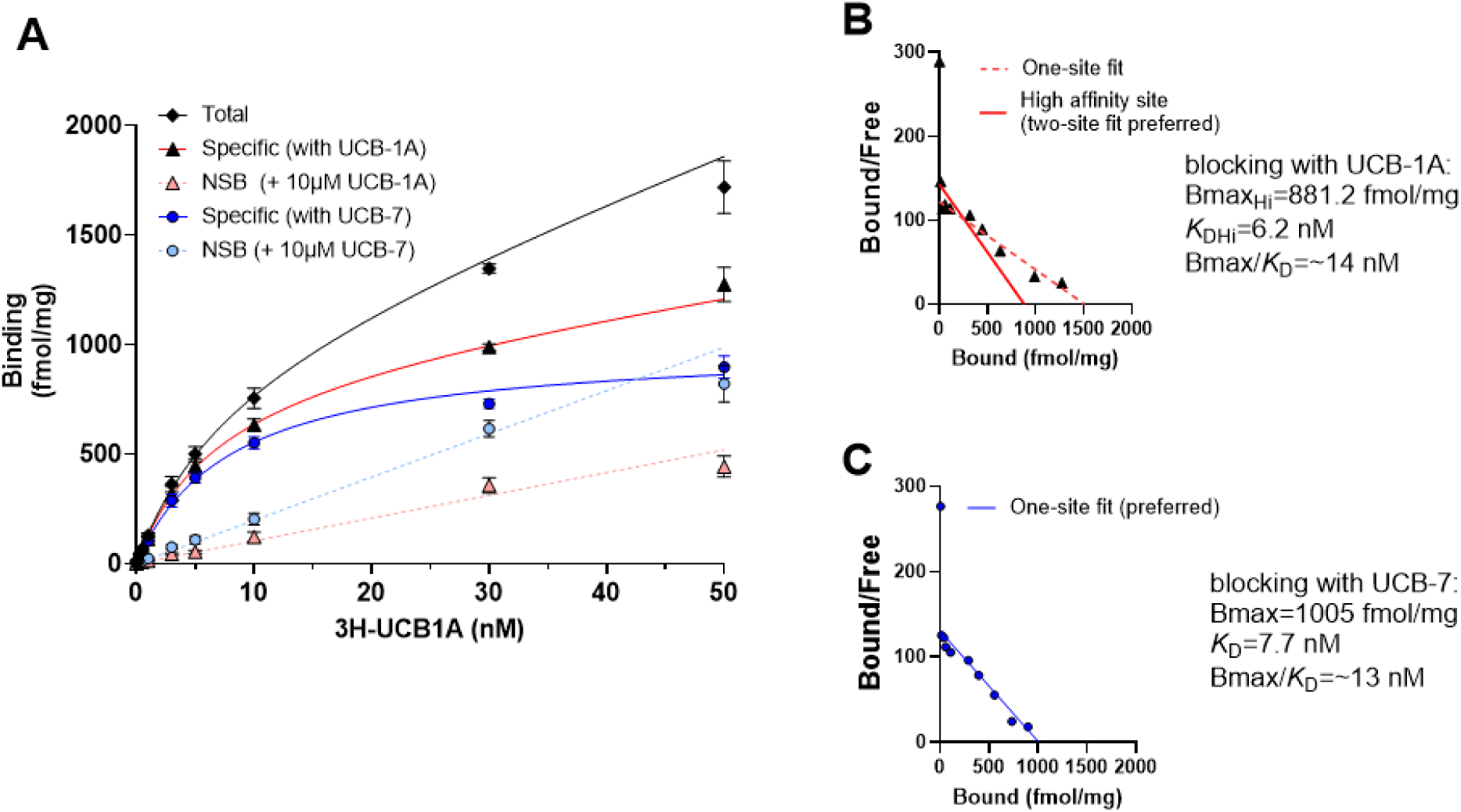
*In vitro* saturation binding studies with [^3^H]UCB-1A in rat brain homogenates from basal ganglia and brainstem. A) Total, non-specific and specific binding of [^3^H]UCB-1A using UCB-7 or unlabeled UCB-1A for blocking. B and C) Scatchard plots for the specific binding in fmol/mg of [^3^H]UCB-1A blocked with UCB-1A (B) or UCB-7 (C), with estimation of B_max_ and *K*_D_.

The estimation of non-specific binding was performed with two SV2C selective blockers (**Table 2**), either UCB-1A (higher affinity for SV2C) or UCB-7 (lower affinity for SV2C). The measured *K*_D_ in brain homogenates from basal ganglia and brainstem was between 6.2 and 7.7 nM with a B_max_/K_D_ ratio between 13 and 14. The ratio of B_max_ between SV2C high- and low-density regions was between 1.5 and 2. In brain homogenates from cortex, hippocampus and cerebellum the B_max_/K_D_ ratio was ∼4.

**Table 2.** In vitro parameters (K_D_ and B_max_) measure in rat brain homogenates.

|  | Non-specific binding estimated using UCB7* |  | Non-specific binding estimated using UCB-1A** |  |
| --- | --- | --- | --- | --- |
|  | BG and brainstem | Ctx, hippocampus and cerebellum | BG and brainstem | Ctx, hippocampus and cerebellum |
| **$K_D$ (nM)** | 6.7 | 17 | 11.2 | 24.9 |
| **$B_{max}$ (fmol/mg)** | 875 | 483 | 1276 | 800 |
| **$B_{max}/K_D$** | 13 | 2.8 | 11.4 | 3.2 |

\*One-site fit was the preferred model for deriving $B_{max}$ and $K_D$
\*\*Two-site fit was the preferred model for deriving $B_{max}$ and $K_D$ for basal ganglia and brainstem (high-affinity site). The values for the low-affinity site were $K_D=1033$ nM and $B_{max}=2182$ fmol/mg, $B_{max}/K_D=0.2$ . The low-affinity site probably corresponds to the binding of [ $^3H$ ]UCB-1A to SV2A (100-fold lower affinity for SV2A vs. SV2C).

### In vitro determination of *K*_D_ of [^3^H]UCB-J on rat brain homogenates

To measure the relative density of SV2C vs. SV2A, saturation binding experiments were performed using [^3^H]UCB-J in rat brain homogenates from the basal ganglia/brainstem and cortex/hippocampus (**fig. S1**). B_max_ and K_D_ values obtained on brain homogenates from the cortex/hippocampus were 10405 fmol/mg protein and 10.3 nM, with a B_max_/K_D_ ratio = 101. B_max_ and K_D_ values obtained on brain homogenates from the basal ganglia/brainstem were 7997 fmol/mg protein and 7.8 nM, with a B_max_/K_D_ ratio = 102. The estimated ratio of B_max SV2A_ / B_max SV2C_ was ∼6.3-to-9.1 in the basal ganglia/brainstem and ∼13-to-21 in the cortex/hippocampus, depending on which compound (UCB7 or UCB1A) was used for determining the non-specific binding (**Table 2**).

To calculate the relative percentage of bound [^3^H]UCB-1A to SV2C and SV2A, we used the experimental B_max_ and K_D_ values. We assumed that [^3^H]UCB-1A has a B_max_/K_D SV2C_ =12 in high-density regions (striatum, pallidus and substantia nigra) and a B_max_/K_D SV2C_ =3 in low-density regions (cortex, hippocampus and cerebellum). Using the B_max_ values for SV2A estimated with [^3^H]UCB-J and assuming a K_D_=400 nM for [^3^H]UCB-1A to SV2A, we estimated for [^3^H]UCB-1A a B_max_/K_D SV2A_ = 2 in the high-density regions and B_max_/K_D SV2A_ = 2.6 in low density regions. Therefore, the relative contribution of [^3^H]UCB-1A binding to SV2C in high-density regions would be [12/(12+2)]=85% and in low-density regions would be [3/(3+2.6)]=54%.

### Autoradiography studies on rat brain tissue sections

Autoradiography studies on rat brain sections (**Fig. 2**) showed that the highest binding of [^3^H]UCB-1A was observed in the pallidum, substantia nigra and brainstem nuclei.

**Figure 2.**
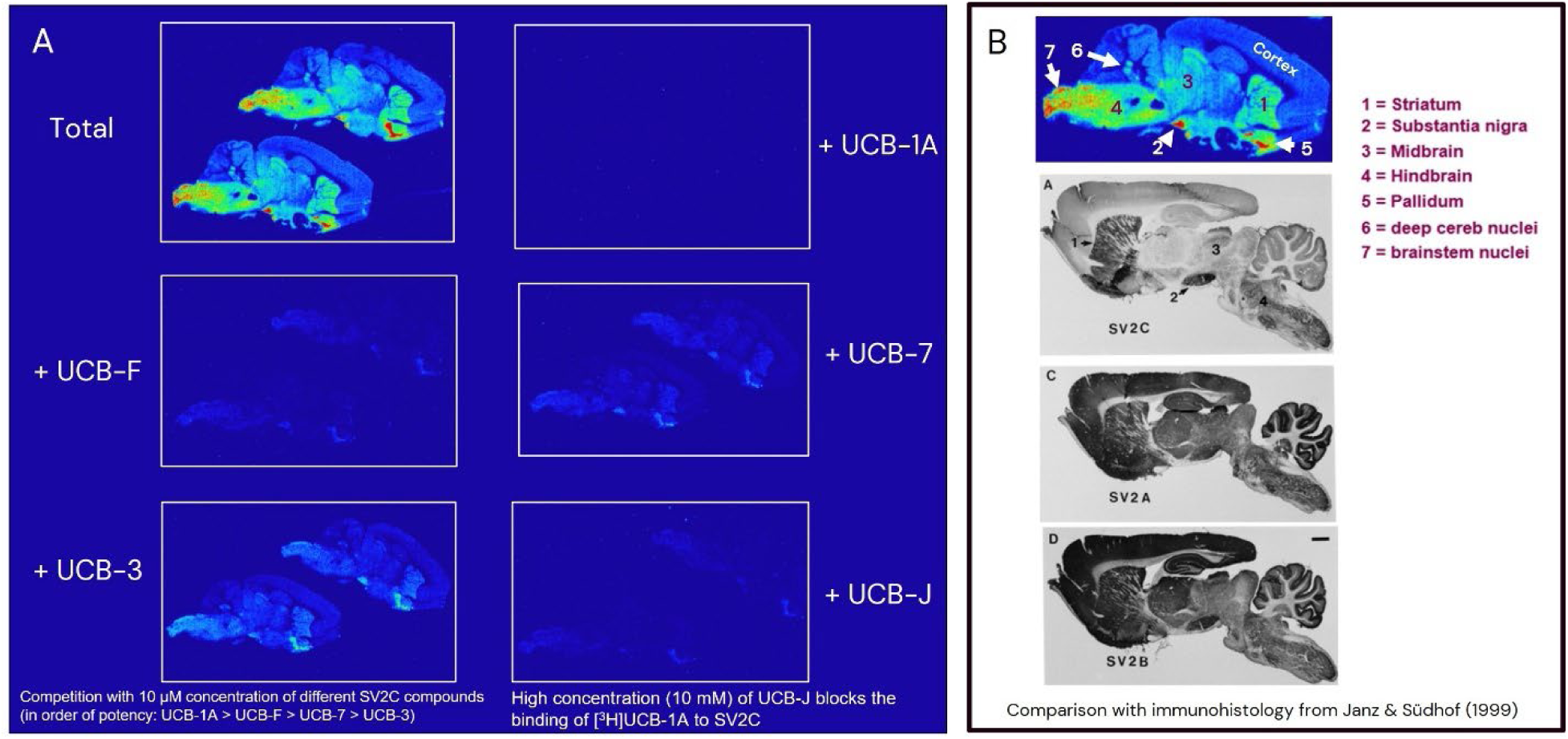
A) Autoradiography study on sagittal rat brain tissue sections showing the total binding of [^3^H]UCB-1A and the binding after competition with 10 μM unlabeled UCB-1A, UCB-F, UCB-7, UCB-3 and UCB-J. B) Total binding of [^3^H]UCB-1A in rat brain showing the distribution consistent with immunohistology data previously reported by Janz and Sudof (*3*).

Lower binding was observed in the striatum, midbrain and deep cerebellar nuclei. The lowest binding was observed in the cortex and cerebellum. The binding of [^3^H]UCB-1A was completely blocked by 10 µM unlabeled UCB-1A. The binding of [^3^H]UCB-1A was also reduced, in order of potency, by 10 µM UCB-F (∼95%), UCB-7 (∼90%), and UCB3 (∼75%) (**fig. S2**). The binding of [^3^H]UCB-1A was reduced by ∼95% also by 10 µM UCB-J. Approximately 50% of the binding of [^3^H]UCB-1A was blocked by 100-to-150 nM UCB-J, which corresponds to the Ki of UCB-J for SV2C (**fig. S3**). Autoradiography studies with increasing concentrations of Levetiracetam provided Ki value of 79 µM in substantia nigra, 49 µM in striatum, 30 µM in cortex (**fig. S4**).

### Autoradiography studies on rat brain tissue sections from 6-hydroxydopamine-lesioned Parkinson model

Autoradiographic studies were conducted on brain sections from rats with unilateral injection of 6-hydroxydopamine (6-OHDA) that were treated with either saline, a single dose of 10 mg/kg L-Dopa, or a chronic treatment with 10 mg/kg L-Dopa for 4 weeks (**Fig. 3**).

**Figure 3.**
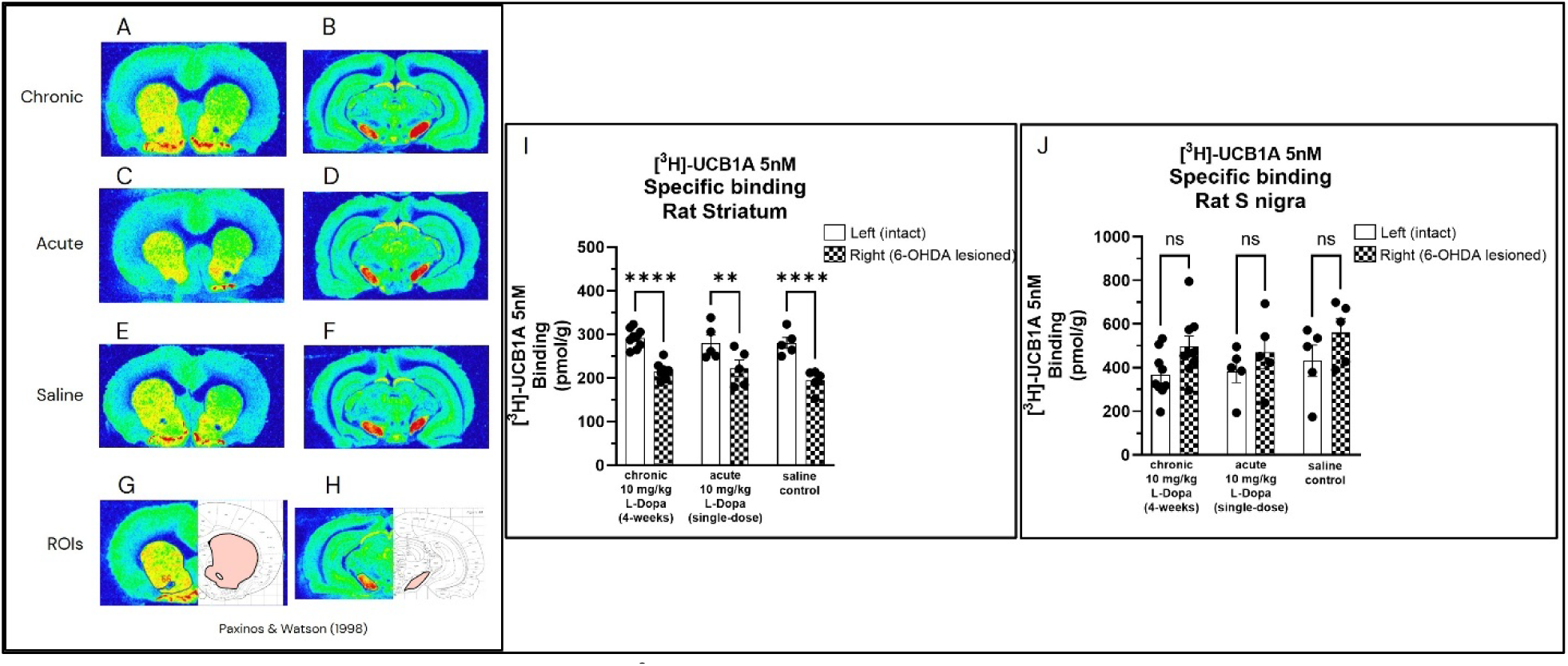
Autoradiography study with [^3^H]UCB-1A on brain tissue sections from rats with unilateral injection of 6-hydroxydopamine (6-OHDA) treated with: A and B) chronic administration of 10 mg/kg L-Dopa for 4 weeks; C and D) acute administration of a single dose of 10 mg/kg L-Dopa; E and F) saline. A, C, E) Coronal sections at the level of the striatum; B, D, F) coronal sections at the level of the substantia nigra. ROIs delineated on G) striatum and H) substantia nigra. Specific binding of [^3^H]UCB-1A in intact and lesioned I) striatum and J) substantia nigra.

In the striatum, significant differences in specific binding of [^3^H]UCB-1A were observed between the lesioned and intact side (two-way ANOVA, F_(1,32)_=59.9, p<0.0001), with no significant differences between saline and L-Dopa treatments. The specific binding of [^3^H]UCB-1A in the lesioned side was 31±6% lower than the specific binding in the intact side in the saline group, 21±6% lower in the acute L-Dopa group and 26±5% lower in the chronic L-Dopa group (**fig. S5**). In the substantia nigra, significant differences in specific binding of [^3^H]UCB-1A were observed between the lesioned and intact side (two-way ANOVA, F_(1,32)_=6.2, p<0.018), with no significant differences between saline and L-Dopa treatments. The specific binding of [^3^H]UCB-1A in the lesioned side was 42±47% higher than the specific binding in the intact side in the saline group, 22±16% higher in the acute L-Dopa group, and 39±32% in the chronic L-Dopa group (**fig. S5**).

### Autoradiography studies in non-human primate and human brain tissue

In non-human primate brain tissue sections, the binding of [^3^H]UCB-1A was highest in the substantia nigra and globus pallidus, followed by the caudate, and putamen, with lowest binding in the thalamus (**Fig. 4**). In the neocortex, moderate binding was observed in the primary motor cortex, sensorimotor cortex and temporal cortex (**Fig. 4**).

**Figure 4.**
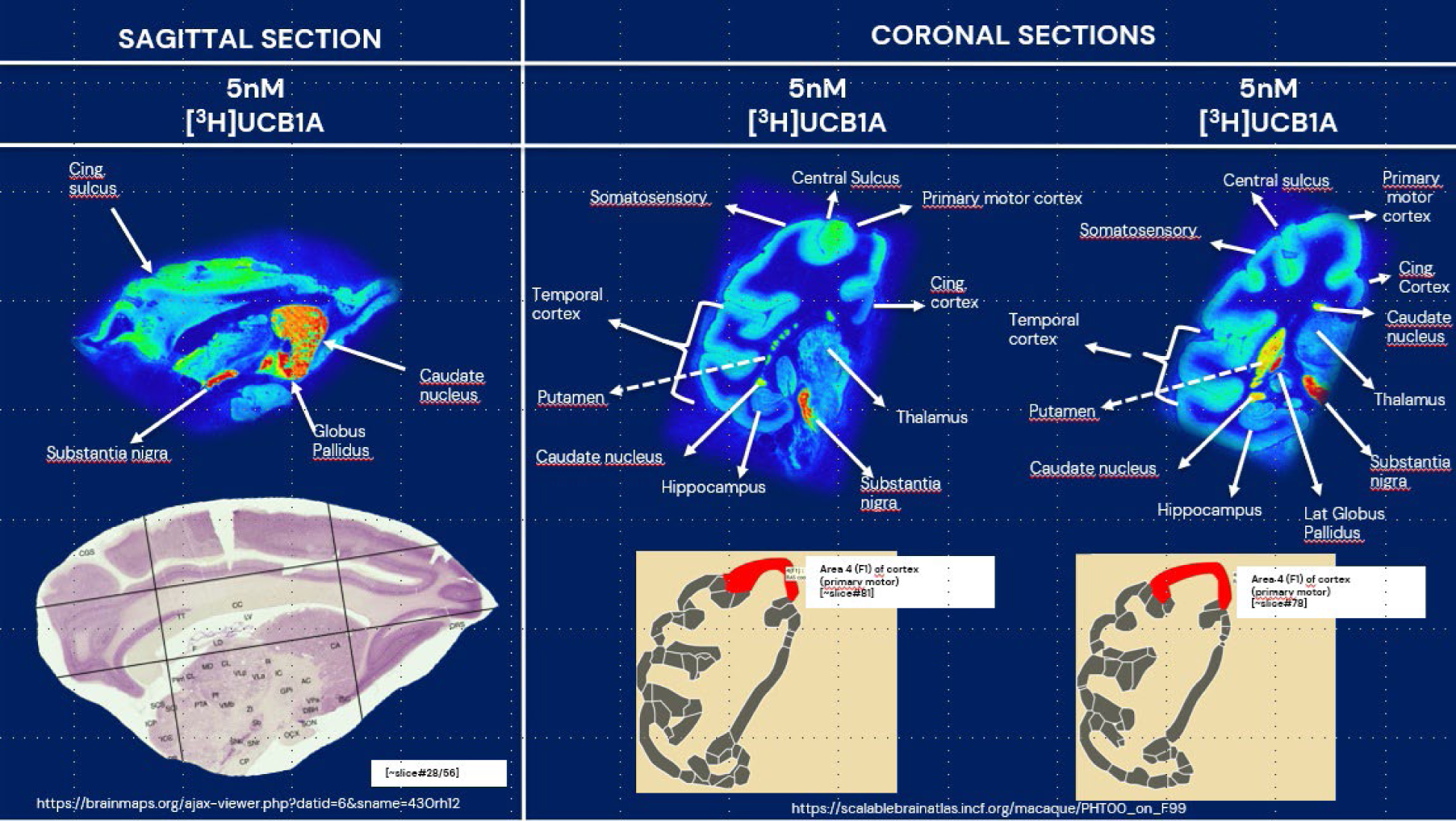
Autoradiography studies with [^3^H]UCB-1A (5 nM) on sagittal and coronal brain tissue sections from three non-human primates.

Lowest binding was observed in the temporal cortex. In the striatum, the distribution of binding of [^3^H]UCB-1A (globus pallidus > caudate > putamen) was very different from the distribution of the binding of the dopamine transporter radioligand [^18^F]FE-PE2I (putamen > caudate > globus pallidus) (**fig. S6**).

Autoradiography studies were conducted with [^3^H]UCB-1A on human brain tissue sections from striatum, globus pallidus, and substantia nigra of five control donors and five Parkinsońs disease donors. The distribution of the binding of [^3^H]UCB-1A in the striatum and substantia nigra was similar as in non-human primates (**Fig. 5**).

**Figure 5.**
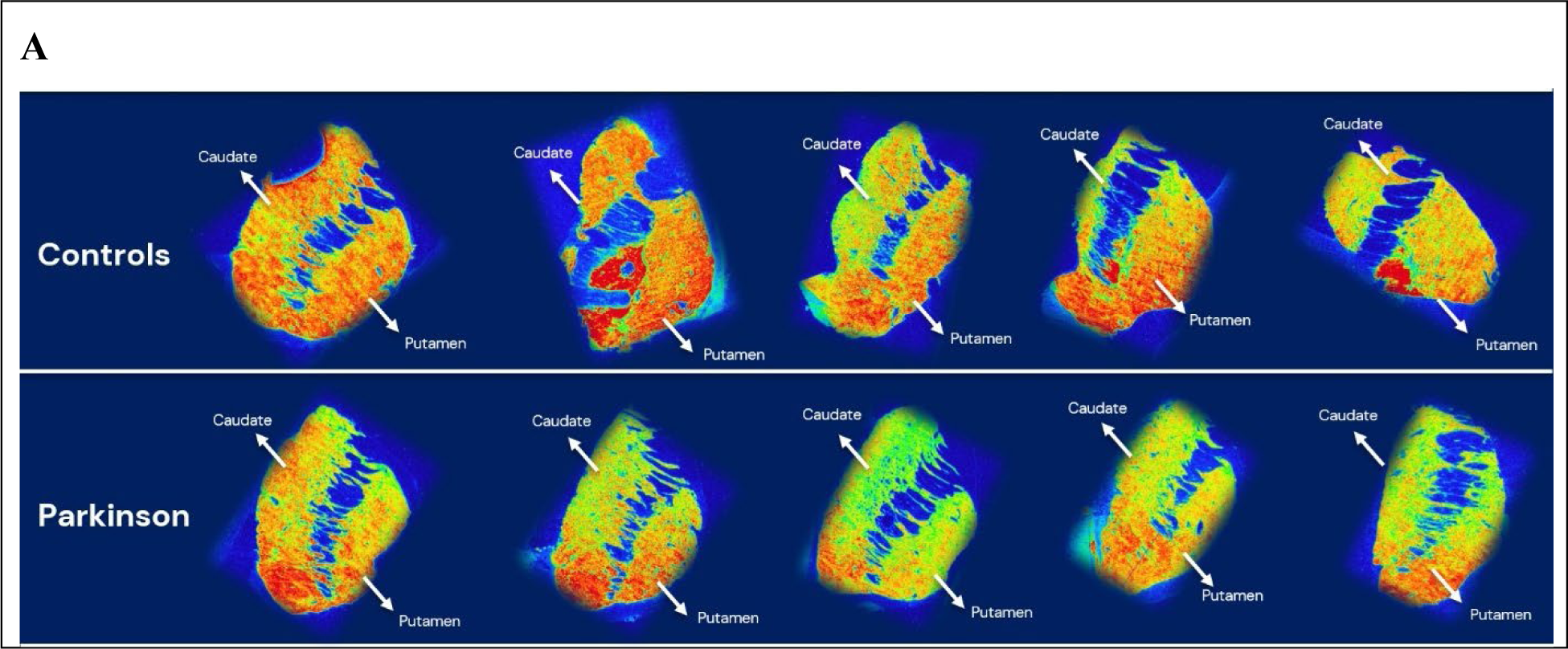
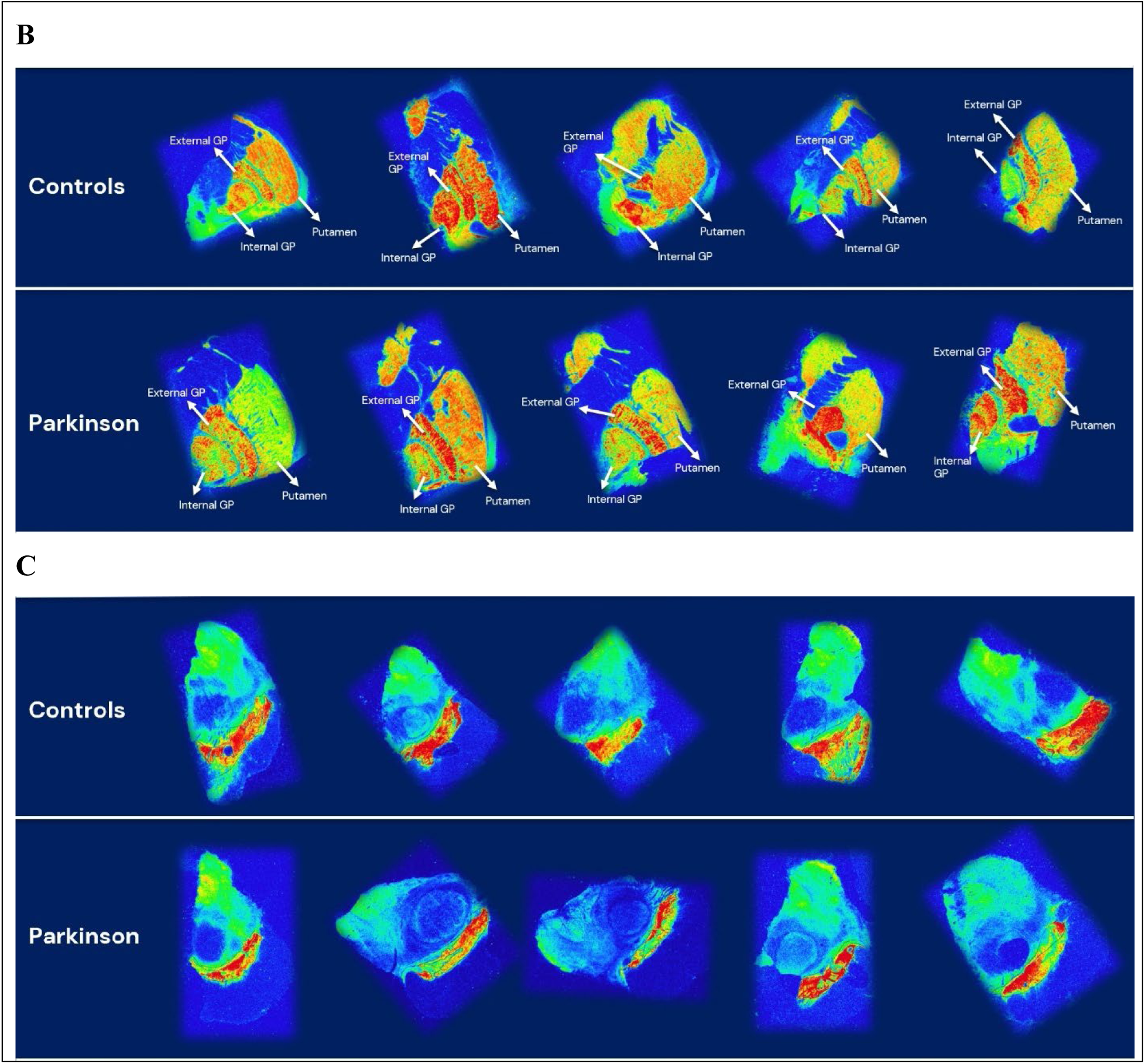
Autoradiography studies with [^3^H]UCB-1A on brain tissue sections from five control donors and five donors with Parkinsońs disease. Coronal sections at the level of A) caudate and putamen and B) putamen and globus pallidus. C) Transaxial sections at the level of the substantia nigra.

The specific binding of [^3^H]UCB-1A in Parkinson donors was lower than in control donors by 12% in the anterior caudate, by 25% in the putamen and by 30% in the posterior putamen (**Fig. 6A**). Higher specific binding of [^3^H]UCB-1A in Parkinson than in control donors was observed in the external (26% higher) and internal globus pallidus (34% higher) (**Fig. 6B**). In the substantia nigra, specific binding of [^3^H]UCB-1A in Parkinsońs disease donors was 8% lower than in controls (**Fig. 6C**). The ratio between external globus pallidus and putamen was 1.25±0.26 in controls and 2.51±0.55 in Parkinsońs disease donors (t-test p=0.01, Coehńs d=1.9, **Fig. 6D**).

**Figure 6.**
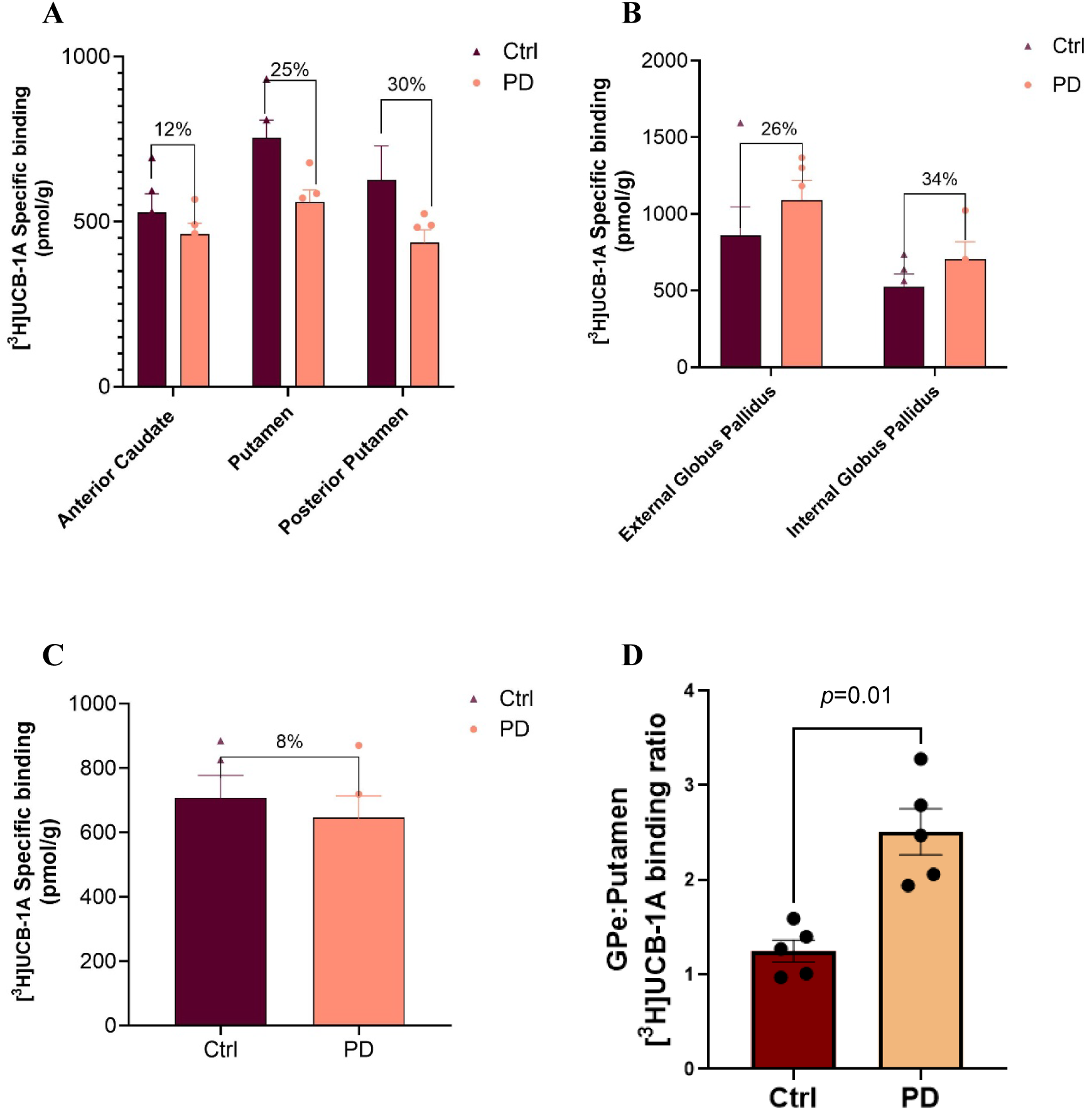
Specific binding of [^3^H]UCB-1A in brain tissue sections of A) striatum, B) globus pallidus, and C) subtantia nigra from control donors (Ctrl) and donors with Parkinsońs disease (PD). D) Ratio between specific binding of [^3^H]UCB-1A in external globus pallidus and putamen in controls (n=5) and Parkinson donors (n=5). Data are presented as mean ± S.E.M.

Autoradiography studies with [^18^F]FE-PE2I on adjacent tissue sections confirmed the known loss of DAT in the striatum and substantia nigra (**fig. S7**). There was no evidence of an effect of age on the specific binding of [^3^H]UCB-1A across subject and brain regions regions.

### Radiolabelling of [^11^C]UCB-1A

The synthesis of [^11^C]UCB-1A was highly reproducible which yielded >1500 MBq of the pure final product following an irradiation of the cyclotron target with a beam current of 35 μA for 20 minutes. Molar activity of the produced radioligand was in a range of 307 ± 129 GBq/µmol (n = 15) at the time of the injection to NHP. The radiochemical purity was >99% and the final product [^11^C]UCB-1A was found to be stable with radiochemical purity more than 98% for up to 60 min.

### In vivo PET imaging in non-human primates

A total of 14 PET measurements with [^11^C]UCB-1A were performed in four non-human primates. Details of the NHPs, injected radioactivity and mass are reported in **Table S2**. Average PET images of [^11^C]UCB-1A (**Fig. 7 and movie S1)**, display the brain distribution of the radioligand, with highest binding in the globus pallidus and substantia nigra, followed by caudate and putamen, medulla and spinal cord, with lower binding in the thalamus. The binding of [^11^C]UCB-1A in the neocortex was lower than in the basal ganglia, with the exception of the sensorimotor cortex. The cerebellum showed the lowest binding among all regions, with the exception of the deep cerebellar nuclei.

**Figure 7.**
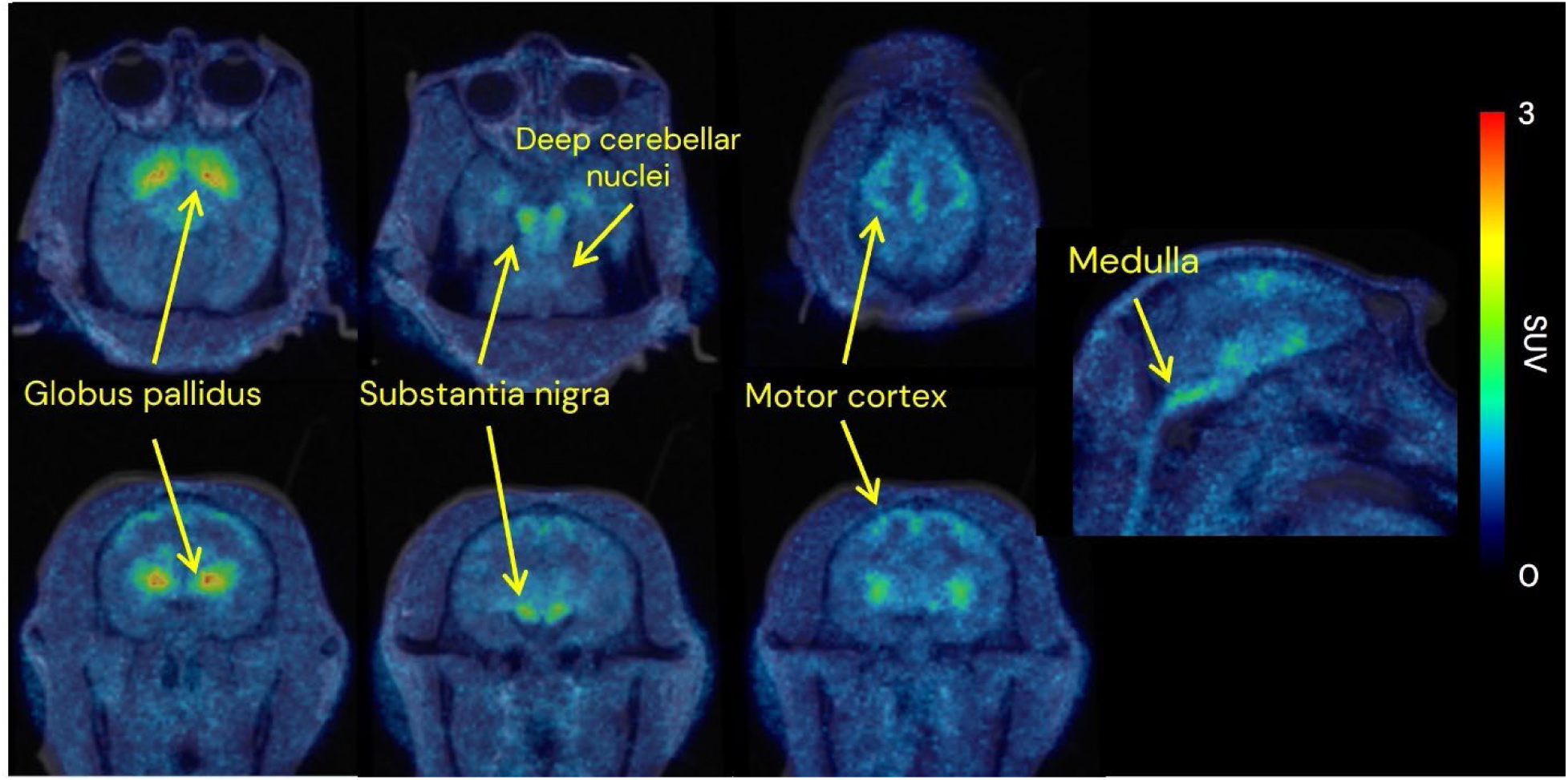
Representative mean images (1-120 min) of [^11^C]UCB-1A in one non-human primate.

Mean (±SEM) time-activity curves of [^11^C]UCB-1A in relevant brain regions are shown in **Fig. 8A**. The t_max_ and C_max_ of [^11^C]UCB-1A for the whole brain were 3 min and 3.5 SUV (**Fig. 8A**).

**Figure 8.**
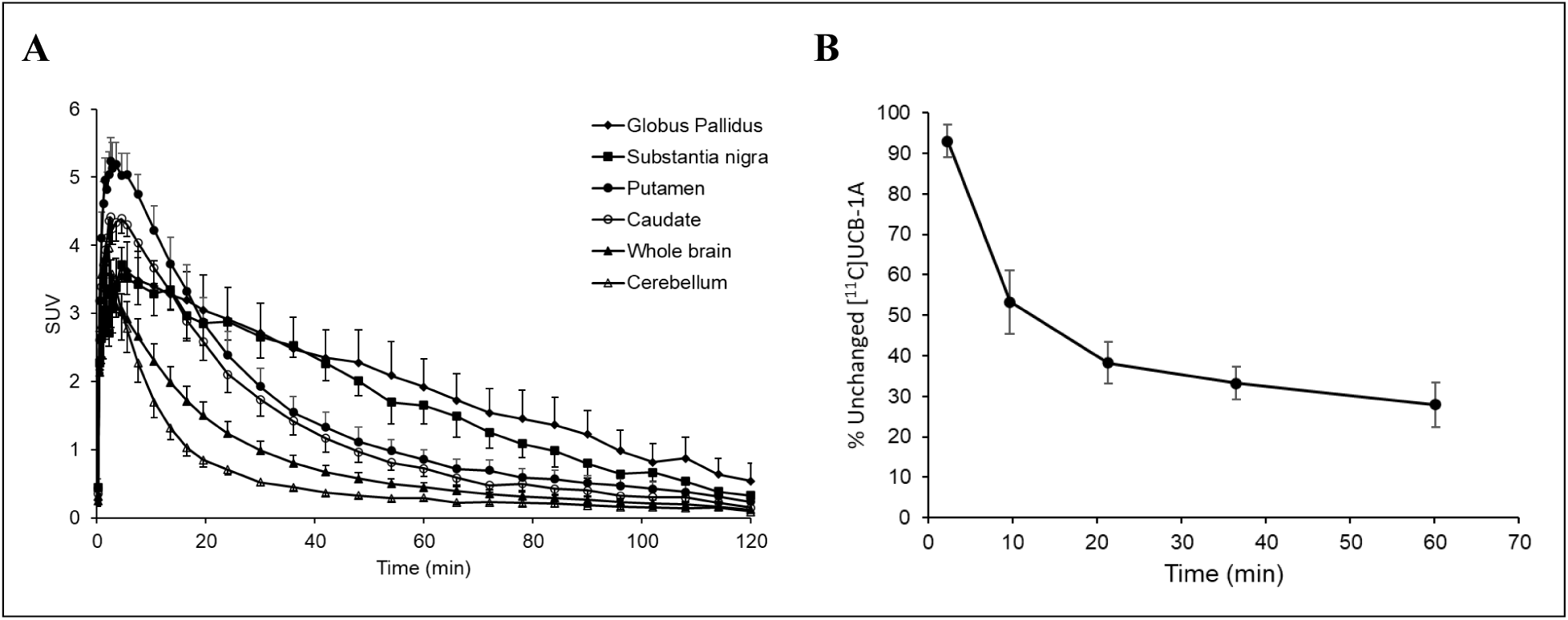
A) Mean (±SEM) time-activity curves and B) mean (±SEM) parent fraction of [^11^C]UCB-1A in 5 non-human primates examined at baseline.

### Radiometabolite analysis, metabolite analysis with mass spectrometry, free fraction in plasma and brain

Radiometabolism of [^11^C]UCB-1A was moderate with approximately 28±5% (mean±SEM) of unchanged radioligand present in plasma at 60 min after radioligand administration (**Fig. 8B**). The HPLC analysis showed that the peak corresponding to [^11^C]UCB-1A had a retention time of ∼5.2 min. Two radiometabolite peaks were also observed, with retention time of ∼3.5 min and ∼4.5 min. These two radiometabolites were more polar than [^11^C]UCB-1A and most likely not brain penetrant (**fig. S8**). Mass spectrometry studies in primary hepatocytes from different species (Caucasian human, Cynomolgus monkey, Sprague Dawley rat and Balb/c mouse) showed that UCB-1A was mainly transformed to doubly oxygenated metabolite M6 (**fig. S8**). This transformation occurs on the pyran and/or the thiadiazole ring. This metabolite was expressed in all non-clinical species and was the most abundant in human, monkey and mouse. The pyran (+N,S from the thiadiazole) was the major site of metabolism. The ethyl group was a secondary site with minor contributions to metabolism (M5) (**fig. S8**). The plasma free fraction (*f*_P_) of UCB-1A was measured in Sprague Dawley rats, cynomolgus and human plasma and the free fraction in brain (fu, brain) was measured in Sprague Dawley rats. The *f*_P_ of UCB-1A in rat was 41.9%, in cynomolgus monkey was 36.8%, in human was 26.3%. the fu, brain in rat was 20.9%. The *f*_P_ measured with ultrafiltration in one NHP (NHP4) was 55%.

### Quantification with 1-TCM, 2-TCM, Logan and MA1

The quantification of the binding of [^11^C]UCB-1A to SV2C was performed with compartmental modelling using 1-TCM and 2-TCM and graphical analysis using Logan and MA1. The preferred model for the analysis of [^11^C]UCB-1A PET data was the 2-TCM (**Fig. 9**). The brain regions with the highest *V*_T_ values were the globus pallidus and substantia nigra, followed by the caudate and putamen, and cerebellum (**Table 3**). Binding of [^11^C]UCB-1A was also observed in other brain regions such as the pons, medulla, spinal cord, sensorimotor cortex, deep cerebellar nuclei. *V*_T_ values estimated with Logan GA and MA1 were in good agreement with those estimated with 2-TCM, although Logan GA and MA1 tended to underestimate *V*_T_, particularly in high-density regions (**Fig. 9**).

**Figure 9.**
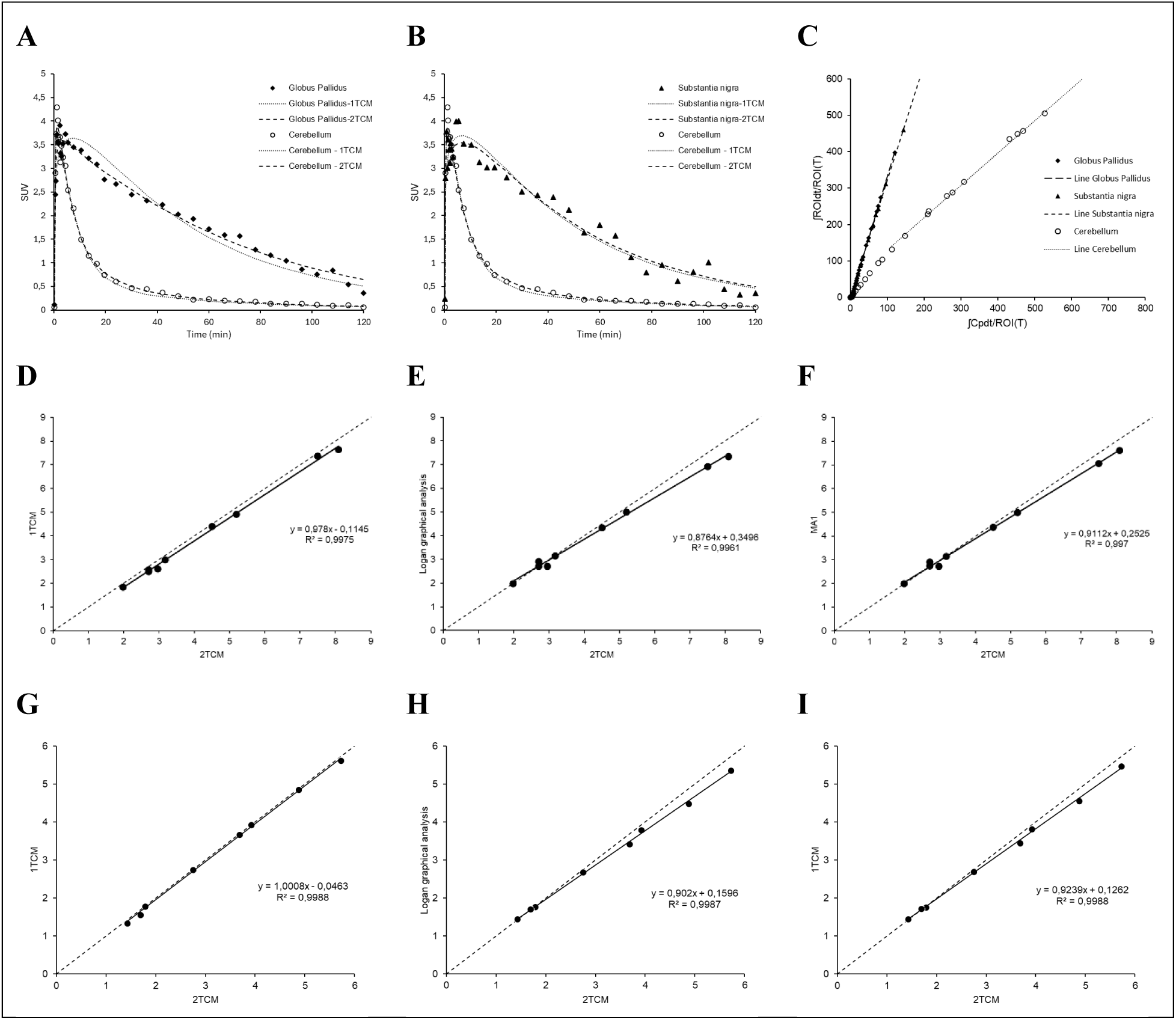
Curve fitting with 1TCM and 2TCM in A) Globus Pallidus, B) Substantia nigra and A,B) Cerebellum for NHP4. C) Logan graphical analysis plots for Globus Pallidus, Substantia nigra and Cerebellum for NHP4. Linear correlations between *V*_T_ estimated with 2TCM and *V*_T_ estimated with D) 1TCM, E) Logan graphical analysis, and F) MA1 in NHP4 and G) 1TCM, H) Logan graphical analysis, and I) MA1 in NHP5.

**Table 3.** Total distribution volume (*V*_T_) or *AUC_ratio_* of [^11^C]UCB-1A in NHPs.

| **NHP4** | **Caudate** | **Putamen** | **Globus pallidus** | **Substantia nigra** | **Cerebellum** |
| --- | --- | --- | --- | --- | --- |
| **Baseline** | 4.5 | 5.2 | 8.1 | 7.5 | 2.0 |
| **10 mg/kg**<br>**Levetiracetam** | 4.1 | 4.5 | 6.9 | 6.5 | 1.6 |
| **Reduction (%)** | 10 | 16 | 17 | 15 | 25 |
| **Baseline** | 4.3 | 4.9 | 7.4 | 6.0 | 1.9 |
| **UCB-1A**<br>**1mg/kg** | 1.8 | 2.1 | 2.1 | 1.8 | 1.6 |
| **Reduction (%)** | 58 | 57 | 71 | 70 | 16 |
| **Test-retest**<br>**variability (%)** | 4.5 | 5.9 | 9 | 22.0 | 5.1 |
| **NHP5** | **Caudate** | **Putamen** | **Globus pallidus** | **Substantia nigra** | **Cerebellum** |
| **Baseline** | 3.7 | 3.9 | 5.7 | 4.9 | 1.4 |
| **30 mg/kg**<br>**Levetiracetam** | 2.7 | 3.0 | 5.0 | 4.0 | 1.1 |
| **Reduction (%)** | 27 | 33 | 12 | 18 | 21 |
| **Baseline\*** | 3.3 | 3.7 | 4.8 | 4.2 | 1.5 |
| **UCB-1A**<br>**1 mg/kg\*** | 1.6 | 1.7 | 1.6 | 1.4 | 1.1 |
| **Reduction (%)** | 52 | 55 | 67 | 67 | 24 |
| **Baseline** | 3.9 | 4.6 | 6.2 | 5.2 | 1.4 |
| **UCB5203**<br>**(0.25 mg/kg)** | 3.9 | 4.3 | 6.9 | 5.3 | 1.3 |
| **Reduction (%)** | 2 | 8 | -10 | -2 | 4 |
| **Test-retest**<br>**variability (%)** | 5.3 | 16.5 | 8.4 | 5.9 | 2.1 |

| **NHP6** | **Caudate** | **Putamen** | **Globus pallidus** | **Substantia nigra** | **Cerebellum** |
| --- | --- | --- | --- | --- | --- |
| **Baseline** | 3.4 | 4.0 | 7.7 | 5.0 | 1.4 |
| **Seletracetam (0.05 mg/kg)** | 3.3 | 4.0 | 8.2 | 5.2 | 1.3 |
| **%Reduction** | 5 | 1 | -6 | -3 | 9 |

\*Arterial cannulation was not possible and instead of $V_T$ the $AUC_{ratio}$ was used as outcome measure. Test-retest variability was calculated from the $V_T$ values obtained at baseline conditions in the same NHP.

The test-retest variability calculated for those NHPs that underwent more than one baseline PET measurement on separate days ranged between 5% to 22% in NHP4 and 5 to 17% in NHP5 (**Table 3**).

### Blocking experiments with Levetiracetam, UCB-1A (SV2C blocker), Seletracetam (SV2A blocker), and UCB5203 (SV2B blocker)

The administration of 10 mg/kg or 30 mg/kg Levetiracetam was associated with an overall decrease of *V*_T_ between 10% and 20% across brain regions (**Fig. 10** and **Table 3**).

**Figure 10.**
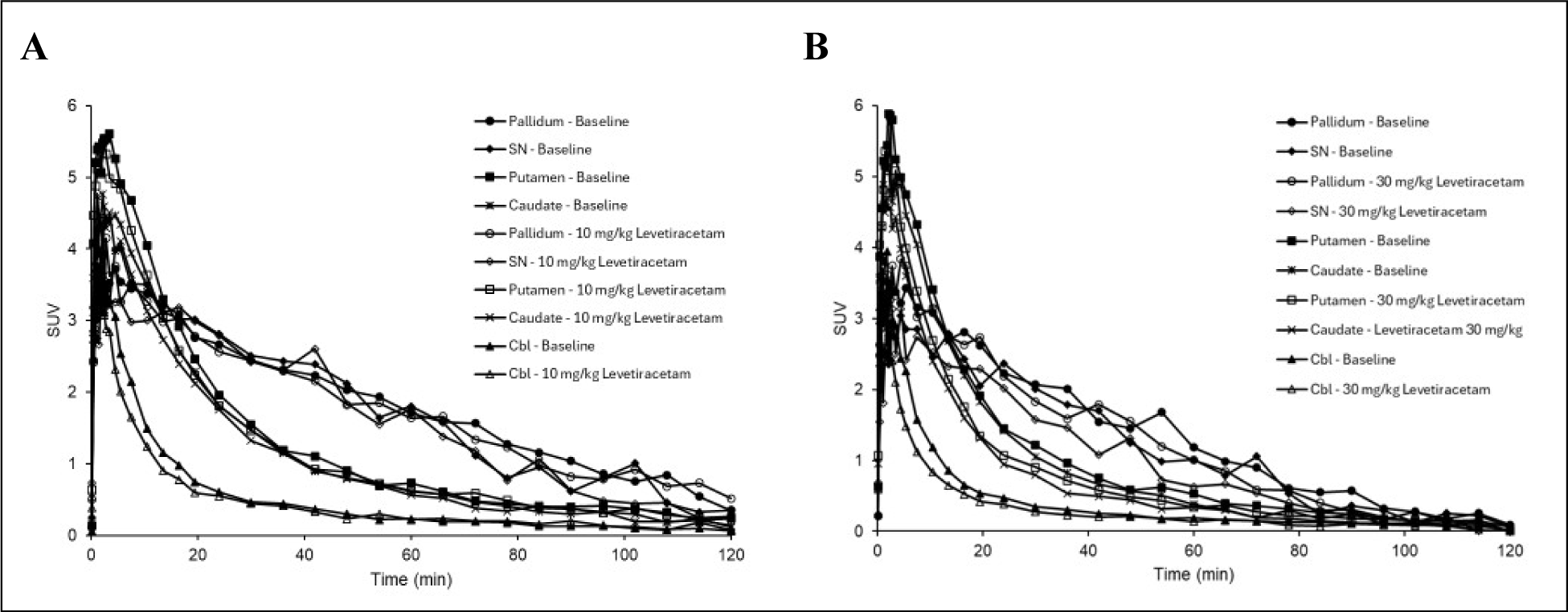
Time activity curves of [^11^C]UCB-1A in caudate, putamen, pallidum, subtantia nigra and cerebellum before and after intravenous administration of Levetiracetam at a dose of A) 10 mg/kg in NHP4 or B) 30 mg/kg in NHP5.

The revised Lassen plot showed a R^2^=0.69 for 10 mg/kg Levetiracetam and R^2^=0.55 for 30 mg/kg Levetiracetam and an estimated occupancy of ∼10% (**fig. S9**). On the other hand, the administration of 1 mg/kg UCB-1A determined a 64% decrease of *V*_T_ and 60% decrease of AUC_ratio_ in high-density regions (caudate, putamen, globus pallidus and substantia nigra), and a 21% decrease of *V*_T_ and a 24% decrease of AUC_ratio_ in cerebellum (**Fig. 11 and Table 3**).

**Figure 11.**
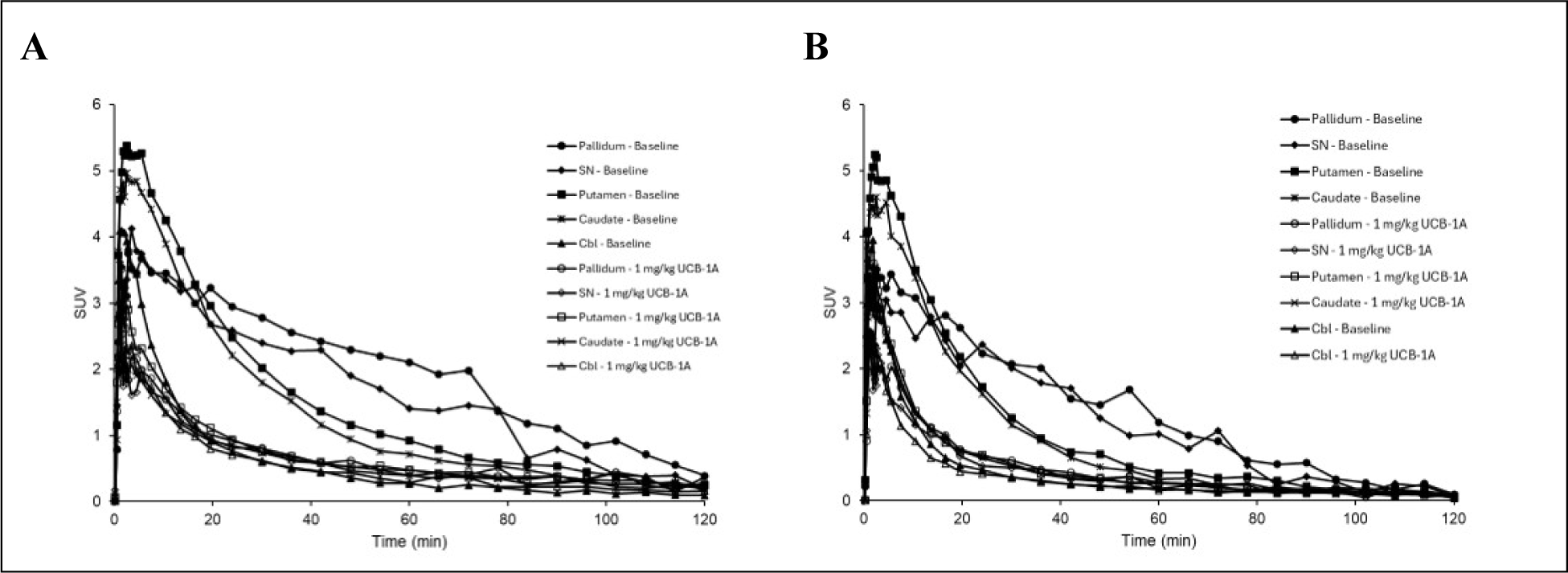
Time-activity curves of [^11^C]UCB-1A in caudate, putamen, pallidum, substantia nigra and cerebellum before and after intravenous administration of 1 mg/kg unlabeled UCB-1A in A) NHP 4 and B) NHP5.

An occupancy of 92% was estimated from the slope of the revised Lassen plot (R^2^=0.997) with an estimated *V*_ND_=1.6 (**fig. S10**). Using the *V*_ND_ value estimated with the revised Lassen plot, we calculated the *BP*_ND_ in all brain regions at baseline and after treatment with either Levetiracetam at the dose of 10 mg/kg or UCB-1A at the dose of 1 mg/kg (**Table S3**). We assumed that at these doses of Levetiracetam would block the binding of [^11^C]UCB-1A to SV2A and UCB-1A would block the binding of [^11^C]UCB-1A to SV2A and SV2C. In high-density regions, after treatment with UCB-1A, the average occupancy in high-density regions was 92%, whereas after treatment with Levetiracetam, the average occupancy in high-density regions was 18%. Therefore, the fraction of [^11^C]UCB-1A bound to SV2C in high density regions would be [(92-18)/92]=80%. The occupancy in the cerebellum with both treatments was >100%, so the fraction of [^11^C]UCB-1A bound to SV2C could not be calculated.

The administration of the selective SV2A compound Seletracetam (0.05 mg/kg) and of the selective SV2B compound UCB5203 (0.25 mg/kg) was not associated with any effects on the binding of [^11^C]UCB-1A to SV2C in any of the brain regions (**Table 3 and fig. S11**). The doses of Seletracetam and UCB5203 were selected based on in vitro binding data (**Table S4**) and predictions of brain exposure from PK data obtained in NHP or extrapolated from rodent PK studies.

## DISCUSSION

The vesicle glycoprotein SV2C is a protein highly expressed in the basal ganglia and involved in the modulation of dopamine release (*2, 14, 15*). The SV2C gene has been found to be associated with Parkinsońs disease and several studies in animal models and brain tissue from Parkinsońs disease donors point towards a link between SV2C and Parkinsońs disease (*7*). We hypothesize that SV2C is a key target for studying synaptopathy in Parkinsońs disease and that PET imaging of SV2C can serve as biomarker for early diagnosis and monitoring of disease progression.

In this study we describe the discovery and the pre-clinical validation of [^11^C]UCB-1A, the first PET radioligand for imaging SV2C. In vitro, UCB-1A displays a *K*_D_ between 6 and 15 nM for SV2C across different species, with a > 100-fold selectivity for SV2C vs. SV2A and SV2B. The binding of [^3^H]UCB-1A in rat brain tissue sections was highest in the pallidum and substantia nigra, followed by the brainstem, the striatum, and the deep cerebellar nuclei, with lowest binding in the cortex and cerebellum. This distribution is consistent with the known expression of SV2C in rats (*2*). In autoradiography studies on brain tissue slices, we found that the binding of [^3^H]UCB-1A was decreased in the striatum of the lesioned side in rats injected with 6-OHDA and in the posterior putamen of Parkinsońs disease donors.

### In vivo PET studies in NHPs

UCB-1A was successfully radiolabelled with ^11^C and PET studies in NHPs showed that [^11^C]UCB-1A displays suitable pharmacokinetic properties, with relatively high brain uptake, reversible kinetics, rapid metabolism without formation of lipophilic radiometabolites. The highest *V*_T_ values were observed in the globus pallidus and substantia nigra, followed by the caudate and putamen, with lowest values in the cerebellum. Blocking studies with unlabeled UCB-1A and selective SV2A and SV2B compounds confirmed in vivo the selectivity of [^11^C]UCB-1A for SV2C. Finally, specific binding of [^11^C]UCB-1A was also observed in the medulla, spinal cord and sensorimotor cortex, all regions involved in motor function. Based on the in vitro and in vivo properties of [^3^H]/[^11^C]UCB-1A and the evidence of reduced binding of [^3^H]UCB-1A in 6-OHDA rats and Parkinsońs disease donors, we believe that [^11^C]UCB-1A is a potential imaging biomarker for Parkinsońs disease.

### Selectivity of [^3^H]/[^11^C]UCB-1A for SV2C

Among all compounds available in the library of UCB compounds, UCB-1A was the most selective SV2C ligand. However, considering the high density of SV2A, some contribution of SV2A to the binding of [^3^H]/[^11^C]UCB-1A cannot be excluded. Considering the relative density of SV2C vs. SV2A in the rat brain and the estimated affinity of UCB-1A for SV2C and SV2A, we have estimated that in high-density regions (striatum, pallidus, substantia nigra and brainstem) a large proportion (∼85%) of UCB-1A is bound to SV2C. A similar value (80%) was obtained by using the in vivo PET data after treatment with UCB-1A and Levetiracetam. Based on the in vitro data, in low-density regions (cortex, hippocampus and cerebellum), ∼54% of UCB-1A is bound to SV2C. In vivo, the contribution of [^11^C]UCB-1A binding to SV2C in cerebellum could not be calculated due to the negligible specific binding of [^11^C]UCB-1A in the cerebellum. In regions with density of SV2C intermediate between basal ganglia and cerebellum (e.g. motor cortex and spinal cord) the proportion of UCB-1A bound to SV2C is expected to be somewhat between 50% and 80%.

### Autoradiography studies on brain slices from 6-OHDA-lesioned rat model and Parkinsońs disease donors

A reduction of specific binding of [^3^H]UCB-1A binding to SV2C in the striatum of 6-OHDA-lesioned rat Parkinsońs disease model and in the putamen of Parkinsońs disease donors was observed. In 6-OHDA-lesioned model, the relatively higher binding of [^3^H]UCB-1A in the substantia nigra of the lesion vs. intact size suggests that the loss of striatal projection induced by the administration of 6-OHDA induces an up-regulation of SV2C in the substantia nigra to compensate for the decreased input through the direct and indirect pathways. The lack of difference in the changes of [^3^H]UCB-1A binding in striatum and substantia nigra between the saline and L-dopa groups suggest that the treatment with L-dopa does not influence the expression of SV2C.

In post-mortem human brain, the reduction of [^3^H]UCB-1A binding in the posterior putamen of Parkinsońs disease donors was observed in combination with a relative increase of [^3^H]UCB-1A binding in the external globus pallidus. This combined effect resulted in an increased ratio between [^3^H]UCB-1A binding in the globus pallidus and [^3^H]UCB-1A binding in the putamen with a very large effect size. The difference of globus pallidus / putamen [^3^H]UCB-1A ratio between Parkinson and control donors might have a diagnostic value in differentiating the two groups if replicated in vivo. In addition, the relative increase of [^3^H]UCB-1A binding in the globus pallidus might reflect an up-regulation of SV2C to compensate for the reduced nigrostriatal input due to the degeneration of the dopaminergic striatal terminals.

In post-mortem human brain tissue, we did not observe a clear difference in the specific binding of [^3^H]UCB-1A between Parkinson and control donors in the substantia nigra. However, the area corresponding to the binding of [^3^H]UCB-1A in the substantia nigra seemed to be smaller in Parkinson than in control donors. Therefore, a potential explanation is that the loss of dopaminergic neurons in the substantia nigra is compensated by an increase of SV2C proteins in the remaining neurons or in the GABAergic terminals. This relative preservation of [^3^H]UCB-1A binding in the substantia nigra in Parkinson donors might reflect similar adaptive changes that were also observed in the substantia nigra of the 6-OHDA rat model. The difference in the magnitude between the effect observed in the 6 OHDA and the effect observed in Parkinsońs disease brain might depend on the type of insult (acute loss vs. chronic degeneration of striatal terminals) and the time span of the natural progression of the human disease.

### Study limitations

This study was designed to develop a PET radioligand for SV2C and included in vitro studies and pre-clinical in vivo studies in NHPs. In vivo data in humans are not yet available. However, a FiH study with [^11^C]UCB-1A is planned in healthy volunteers and participants with Parkinsońs disease. This study will evaluate the suitability of [^11^C]UCB-1A as PET radioligand for imaging SV2C in human subjects and whether differences in SV2C availability observed in vitro between Parkinsońs disease patients and controls will be confirmed in vivo. The evidence of a blocking effect of UCB-1A in the cerebellum suggest that this region cannot be used as reference tissue for simplified quantification of [^11^C]UCB-1A binding to SV2C. In analogy with [^11^C]UCB-J PET, the quantification will require arterial cannulation for human studies. However, the cerebellum displays the lowest *V*_T_ and negligible specific binding. Therefore, if the cerebellum is not affected by the disease (e.g. in case of Parkinsońs disease) it could be likely used as (pseudo)reference region, which will enable simplified quantification in future studies.

### Clinical use of [^11^C]UCB-1A as synaptic imaging marker in neurodegenerative disorders

[^11^C]UCB-1A is a selective PET radioligand with suitable pharmacokinetic properties for imaging and quantification of SV2C in the brain. Our in vitro studies showed a reduction of the specific binding of [^11^C]UCB-1A in the striatum of 6-OHDA model and of donors with Parkinsońs disease. Taken together, our findings suggest that [^11^C]UCB-1A is a promising synaptic imaging marker for the study of patients with Parkinsońs disease. We believe that [^11^C]UCB-1A can serve as diagnostic imaging agent and as biomarker for evaluation of disease progression and treatment effects. Finally, the distribution of [^11^C]UCB-1A in brain regions primarily involved in the cortico-spinal motor pathway suggest the potential utility of the radioligand as imaging biomarker also in other neurodegenerative disorders, such as Amyotrophic Lateral Sclerosis.

## MATERIALS AND METHODS

### Autoradiography experiments on rat, NHP and human brain tissue

#### Animals

Naïve, adult Wistar rats were bred at the animal facility in the Department of Comparative Medicine at the Karolinska Institute and euthanized by placing them in a CO2 saturated chamber followed by decapitation. Brains were quickly removed, and snap frozen (whole, or dissected into regions) in isopentane (Sigma-Aldrich, St. Louis, MO, USA) cooled in dry ice, and subsequently stored at -80° C.

Studies using NHP brain tissue were approved by the Animal Ethics Committee of the Swedish Animal Welfare Agency (registration no. 4820/06-600 and 399/08). Sections from the brain of 1 adult female cynomolgus monkey (Macaca Fascicularis) were used in this study.

#### Post-mortem Human Tissues

The human *post-mortem* brain sections were obtained from the “Syndey Brain Bank” and it’s use in this study was approved by the Swedish Ethical Research Authority. The *post-mortem* samples consisted of n=5 PD donors (4 females, 1 male, 79±2 y) and n=5 non-dementia control donors (all female, 93±4 y, disease duration 7 to 26 y). From each donor, tissue blocks containing the following regions were used: 1) anterior caudate and putamen, 2) posterior putamen and globus pallidus, and 3) substantia nigra.

#### Autoradiography workflow

The sections were first thawed at room temperature and then pre-incubated for 20 minutes with 50 mM Tris HCl Binding buffer, pH = 7.4 (120mM NaCl, 5mM KCl, 2mM CaCl2, 1mM MgCl2), followed by 1-hour incubation with [^3^H]UCB-1A (5 nM) alone (Total) or co-incubated in presence of UCB-1A (10 µM) to determine the NSB. Then, the slides were washed two times with cold (4°C) 50 mM Tris washing buffer, pH=7.4 and then dipped in water. At the end of the last wash steps, the sides were dried with paper and placed on a plate heater until they looked completely dried. The slides were then placed in cassette together with tritium standards on glass slides (American Radiolabeled Chemicals Inc., St. Louis, MO, USA) and phosphor imaging plates (Fujifilm Plate BAS-TR2025, Fujifilm, Tokyo, Japan) were placed over the slides to expose. The cassette was placed in a container shielded from background and cosmic radiation. The exposure time was 90 hours and then the scanning was performed using an Amersham Typhoon FLA-9500 phosphor imaging scanner (Cytiva, Marlborough, MA, USA).

### Preparation of brain sections from striatum and substantia nigra of 6-hydroxydopamine lesioned rats

Adult Sprague–Dawley rats were used. Experiments were performed in agreement with the European Communities Council (86/609/EEC) and were approved by the ethical committee at Karolinska Institute (N282/06). Rats were anesthetized with 80 mg/kg ketamine (i.p.; Parke-Davis) and 5 mg/kg xylazine (i.p.; Bayer), pretreated with 25 mg/kg desipramine (i.p.; Sigma–Aldrich) and 5 mg/kg pargyline (i.p.; Sigma–Aldrich), placed in a stereotaxic frame, and injected with 12.5 μg of 6-OHDA into the right median forebrain bundle (MFB). The coordinates for injection were AP, −2.8 mm; ML, −2.0 mm; and DV, −9.0 mm relative to bregma and the dural surface. Two weeks after unilateral 6-OHDA lesioning, rats were administered 1 mg/kg apomorphine (i.p; Sigma–Aldrich). Only rats rotating >100 turns per 30 min were included in further experiments. Four weeks after surgery, rats were treated with saline or 10/7.5 mg/kg L-Dopa/benserazide (i.p., once daily) for one day or 28 days. Animals were killed 1 h after the last injection. Brains were quickly removed, and snap frozen in isopentane (Sigma-Aldrich, St. Louis, MO, USA) cooled in dry ice, and subsequently stored at -80° C. Frozen rat brains were cut with a cryotome (Leica Microsystems AB, Kista, Sweden) at 12 μm thickness.

### In-silico modeling of the binding of UCB-1A to SV2 proteins

A computational approach with molecular dynamics simulations was used to study the binding of UCB-1A to SV2 proteins. The crystal structure of human SV2A in complex with a small-molecule ligand (PDB ID: 8UO9) was used as the initial template. Complexes were embedded in a palmitoyl-oleoyl-phosphatidylcholine (POPC) bilayer using the Schrödinger System Builder, with explicit TIP3P water molecules added for solvation. Neutralizing counterions (Na⁺, Cl⁻) were included to approximate the physiological ionic strength. Ten ns molecular dynamics was applied for the SV2A–UCB1A model employing the OPLS4 force fields. Homology models were also generated for SV2C, using the relaxed SV2A–UCB1A structure as a template. The modeled SV2C–UCB1A complexes were each refined by a 10 ns MD relaxation to stabilize side-chain conformations and optimize ligand positioning. Temperature effects were simulated under two conditions: low temperature - 277 K (4 °C), and high temperature - 310 K (37 °C), representing physiological body temperature. Production simulations were conducted for 100 ns per trajectory, with 10 independent replicas performed for each temperature condition to enhance statistical robustness. A combination of Schrödinger simulation analysis tools and in-house Python scripts were applied for the trajectory analyses.

### Radiolabelling of [^11^C]UCB-1A

The precursors UCB-1 (3-(3-ethyl-1H-pyrazol-5-yl)-6-(tetrahydro-2H-pyran-4-yl)-[1,2,4]triazolo [3,4-b][1,3,4]thiadiazole) and the non-radioactive reference standard UCB-1A (3-(3-ethyl-1-methyl-1H-pyrazol-5-yl)-6-(tetrahydro-2H-pyran-4-yl)-[1,2,4]triazolo[3,4-b][1,3,4] thiadiazole) were synthesized by Aurigene Pharmaceutical Services Limited, Bangalore 560100, India. [^11^C]Methyltriflate ([^11^C]CH3OTff), was synthesized according to the previously published method (*25, 26*). [^11^C]UCB-1A was made by trapping [^11^C]CH3OTf with the amine precursor (UCB-1, 1.0–1.5 mg) in acetone (400 μL), heating 50 °C for 60 s. The crude was diluted (2 mL water) and purified by HPLC. The [^11^C]UCB-1A fraction eluted at tR 9–10 min and was collected into sterile water (50 mL). The pooled product was purified on a preconditioned Oasis HLB 3 cc SepPak cartridge, washed with sterile water (10 mL), and eluted with ethanol (1 mL) into a sterile vial containing sterile saline (9 mL). The final product was sterile filtered through a 0.22 μm Millipore Millex GV unit. The total time for radiosynthesis including HPLC purification, SPE isolation and formulation was about 35 min.

### PET studies in non-human primates

The study adhered to ethical standards and received approval from the Animal Ethics Committee in Stockholm, administered by the Swedish Board of Agriculture (10367-2019 and 9878-2024). The study also complied with the guidelines outlined in "Guidelines for Planning, Conducting, and Documenting Experimental Research" (Dnr 4820/06-600) of Karolinska Institutet. The non-human primates (NHPs) used in this study were housed at the Astrid Fagraeus Laboratory, Comparative Medicine, in Solna, Sweden. Five male cynomolgus monkeys (*Macaca fascicularis*) weighing 6 to 10 kg were selected for participation.

At the Astrid Fagraeus Laboratory, each non-human primate (NHP) received ketamine hydrochloride at 10 mg kg-1 by intramuscular injection to induce anaesthesia. Endotracheal intubation-maintained anaesthesia with a blend of sevoflurane, oxygen and medical air. Body temperature was maintained using a Bair Hugger model 505 warming unit (Arizant Healthcare, MN) and monitored with an esophageal thermometer. Heart rate, blood pressure, respiratory rate and SpO2 were recorded on a bedside monitor, and saline dripped continuously to correct volume losses. PET measurements were performed using the LFER 150 PET/CT system (Mediso Ltd.) (*27*). The head of the monkey was immobilized using the specifically designed NHP bed of the LFER 150. An arterial catheter was inserted in one of the iliac arteries under ultrasound guidance. [^11^C]UCB-1A was administered as bolus, followed by the administration of 2 mL NaCl.

PET data were acquired in list mode for 123 min and reconstructed with a series of frames of increasing duration (9 x 20 sec, 3 x 60 sec, 5 x 180 sec, 17 x 360 sec). The reconstruction was performed using the Tera-Tomo 3D algorithm (with 9 iterations and 5 subsets), including detector modeling with attenuation and scatter correction based on a 5-component material map. Volumes of interest (VOIs) were delineated on an MR image of the animal, which was manually coregistered with the average PET image using the FUSION tool in PMOD software (version 4.2; Bruker). Selected VOIs were caudate, putamen, globus pallidus, substantia nigra, and cerebellum. Radioactivity concentration was expressed as SUV. Arterial blood samples were collected for the first 3 min using an automated blood sampling system (Veenstra, Comecer) at a speed of 3 ml/min. Manual samples were collected at 1.5, 2.5, 3 and 120 min after injection for measurement of blood and plasma radioactivity, and at 2, 5, 15, 30, 60, 75, and 90 min for radioactivity measurements and radiometabolite analysis.

PET quantification was performed with the PKIN tool of PMOD, using one-tissue, two-tissue compartment models (1-TCM and 2-TCM), Logan graphical analysis and MA1. Blocking studies were performed with the IV administration by 5-min infusion of Levetiracetam (10 mg/kg and 30 mg/kg), UCB-1A (1 mg/kg), seletracetam (0.05 mg/kg), UCB5203 (0.25 mg/kg), 20 min before radioligand administration.

### Radiometabolite analysis

Radiometabolite analysis of [^11^C]UCB-1A was performed using a radio-HPLC system consisting of an interface module (D-7000; Hitachi, Tokyo, Japan), an L-7100 pump (Hitachi), and a manual injector (Model 7125, Rheodyne, Cotati, CA, USA) equipped with a 5.0-mL sample loop. Ultraviolet detection was carried out at 254 nm using an L-7400 UV detector (Hitachi), connected in series with a Packard 150TR radioactivity detector housed in 50-mm lead shielding and equipped with a 550-µL flow cell. Chromatographic separation was achieved on an ACE 5 µm C18 HL column (250 × 10.0 mm). The mobile phase consisted of acetonitrile (A) and 0.1 M ammonium phosphate buffer (pH 7.0) (B), delivered at a flow rate of 5.0 mL/min under gradient conditions as follows: 0–4.0 min, 40:60 to 75:25 (A/B, v/v); 4.0–6.0 min, 75:25 (A/B, v/v); 6.0–6.1 min, 75:25 to 40:60 (A/B, v/v).

## Acknowledgments

The authors thank the staff of Brain Molecular Imaging Centre for the conduction of the PET experiments in non-human primates. We also thank the families and donors for providing consent to brain donation for research.

## Author contributions

Conceptualization: AV, SN, CH, HÅ

Methodology: AV, SN, YKM, AFM, R, CH, HÅ, VS, YZ, PS

Investigation: AV, SN, YKM, AFM, R, CH, HÅ, VS, JM, YZ, PS

Visualization: AV, SN, JM, R

Funding acquisition: AV, CH, HÅ

Project administration: AV, SN

Supervision: AV, SN, CH, HÅ, JM, PS

Writing – original draft: AV, SN, VS, AFM, R, JM, YZ

Writing – review & editing: AV, SN, CH, HÅ, YKM, AFM, VS, JM, YZ, PS

## Data and materials availability

Proprietary compounds of UCB for in vitro and in vivo experiments have been provided to KI under research collaboration and materials transfer agreement. Data can available upon reasonable request to the corresponding author.

