## Supplementary Material for "Discovery and characterization of UCB-1A: the first PET radioligand for imaging synaptic vesicle glycoprotein 2C"

**Affiliations:**

### List of Supplementary Materials

Fig. S1 to Fig. S11. Tabls S1 to Table S4. Movie S1.

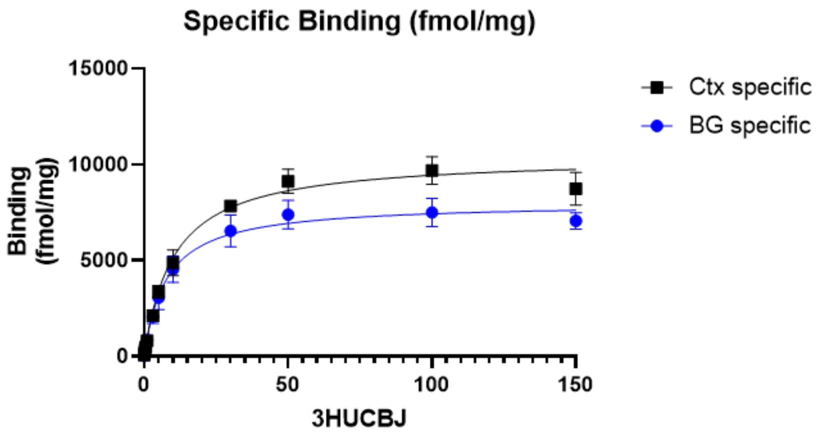


| | Ctx specific | BG specific |
|---|---|---|
| One site -- Specific binding | | |
| Best-fit values | | |
| Bmax | 10405 | 7997 |
| Kd | 10.31 | 7.749 |

**Figure S1**. In vitro binding assay with [$^3$H]UCB-J in rat brain homogenates from cortex and basal ganglia.

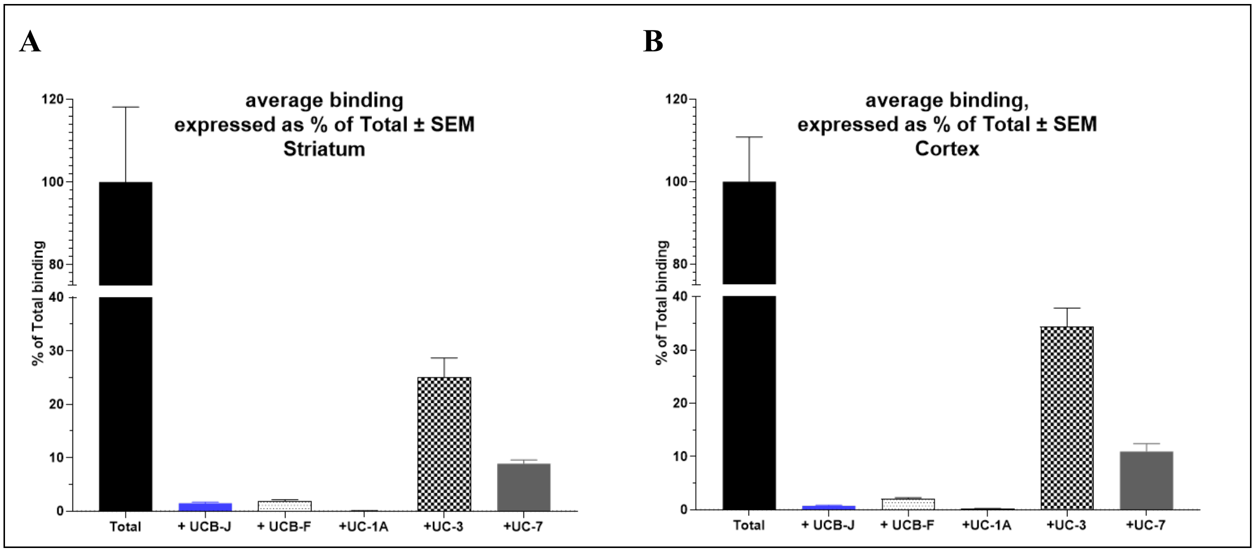


**Figure S2**. Quantification of autoradiography studies with [$^3$H]UCB-1A (5 nM) in rat brain homogenates from A) striatum and B) cortex. Each value is the average (+/- SEM) of n=3 rats, each of which is in turn averaged from 4 technical replicates.

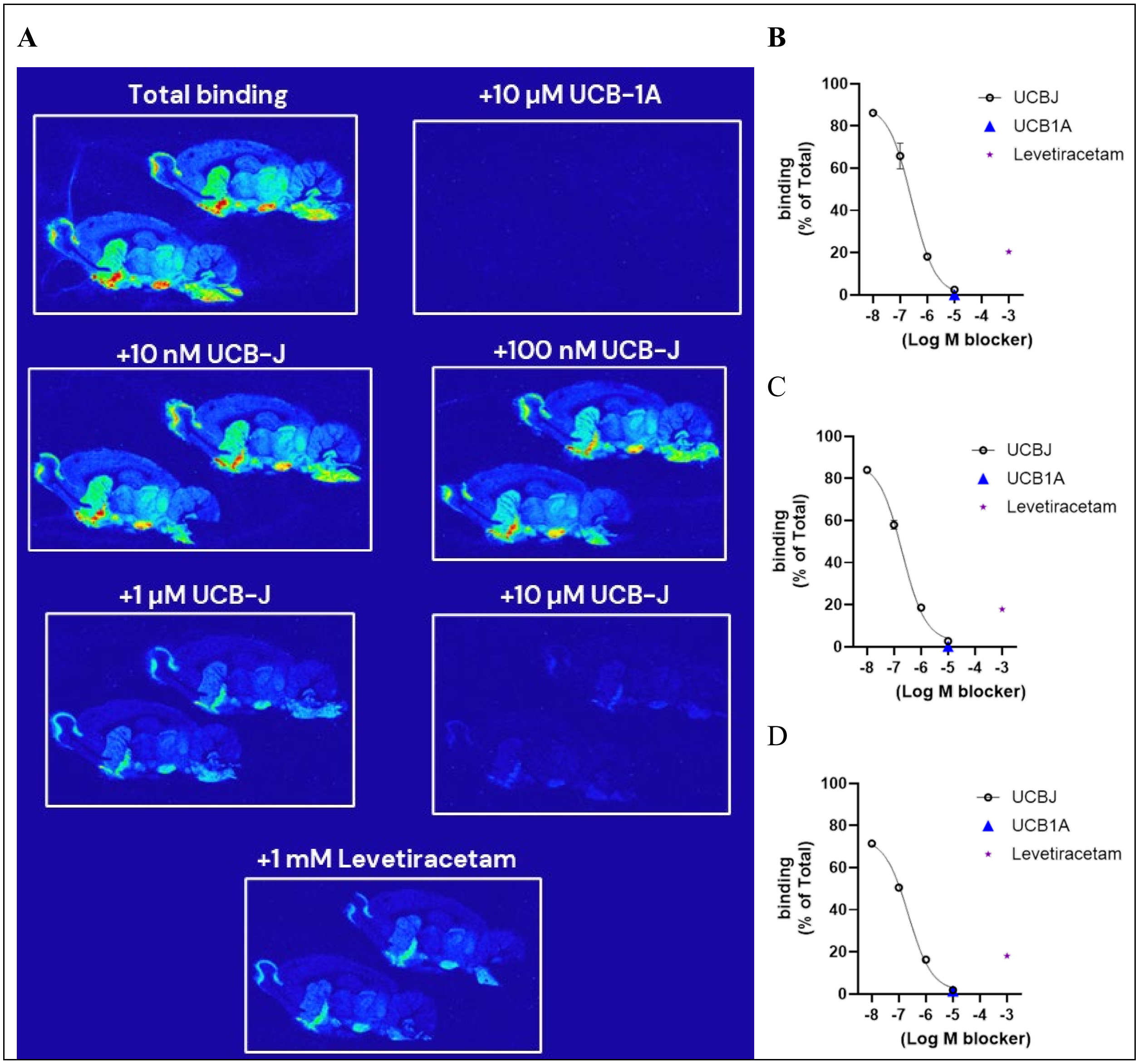


**Figure S3**. A) Autoradiography studies with [$^3$H]UCB-1A (5 nM) in rat brain sections with UCB-1A (10 µM), increasing concentration of UCB-J (from 10 nM to 10 µM) and Levetiracetam (1 mM). Percent of total binding measured in B) substantia nigra, C) striatum, and D) cortex. Each plotted value is the average (+/- SEM) of n=3 rats, each of which is in turn averaged from 4 technical replicates.

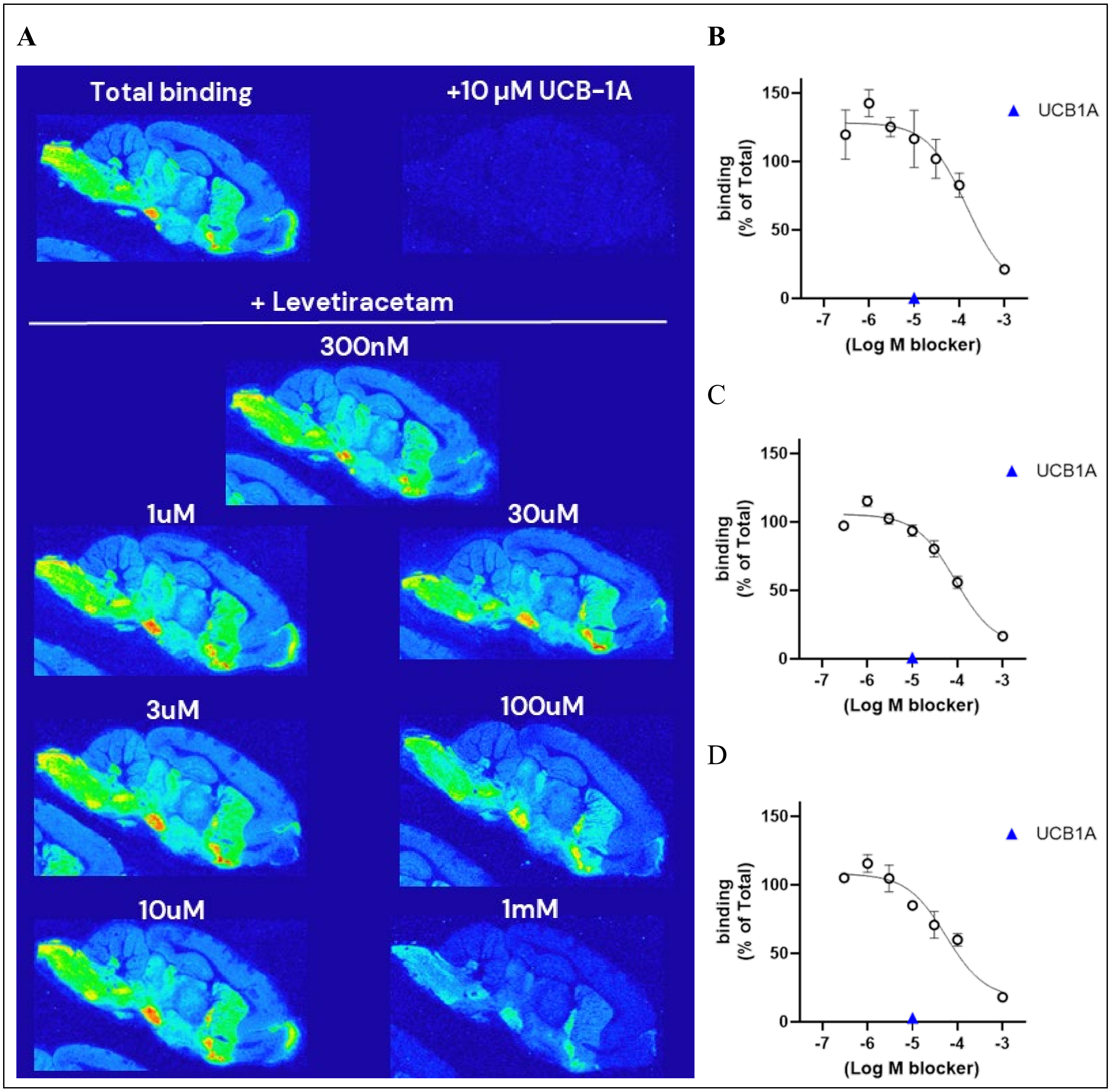


**Figure S4**. A) Autoradiography studies with [$^3$H]UCB-1A in rat brain sections with UCB-1A (10 µM), and increasing concentration of Levetiracetam (from 300 nM to 1 mM). Percent of total binding measured in B) substantia nigra, C) striatum, and D) cortex. Each plotted value is the average (+/- SEM) of n=3 rats, each of which is in turn averaged from at least 2 technical replicates.

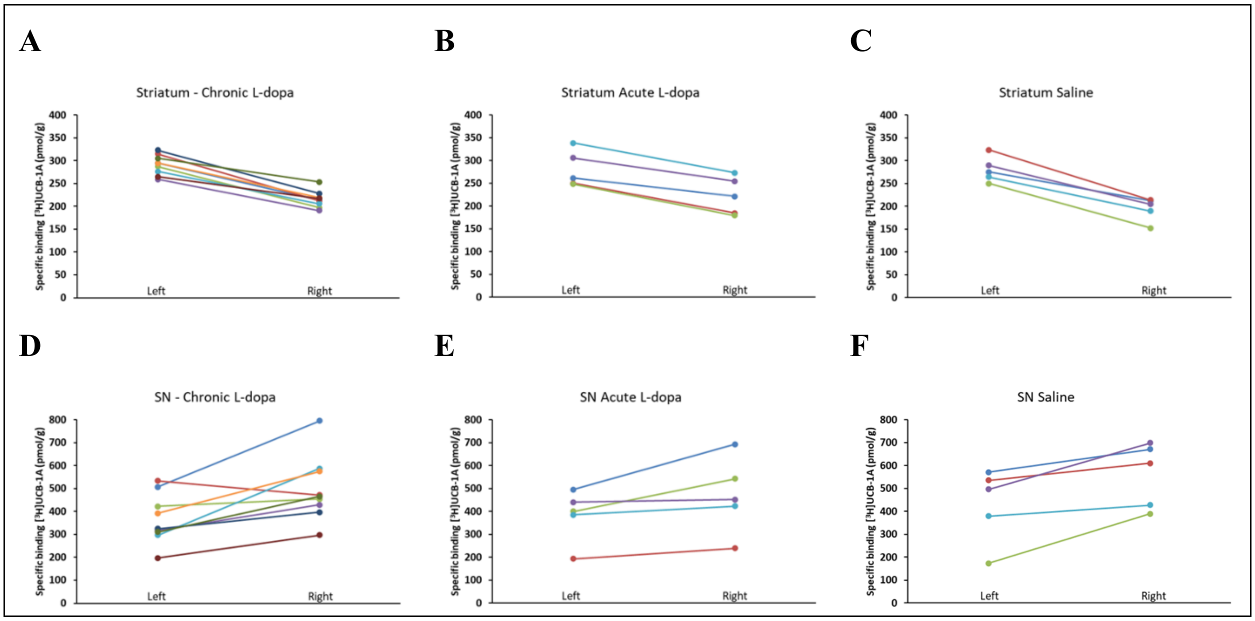


**Figure S5**. Specific binding of [$^{3}$H]UCB-1A in 6-OHDA rats in A, B, and C) the striatum and D, E, F) substantia nigra of the lesioned (right) and non-lesioned (left) side. Animals were treated with A, D) chronic L-dopa; B, E) acute L-dopa, and C, F) saline.

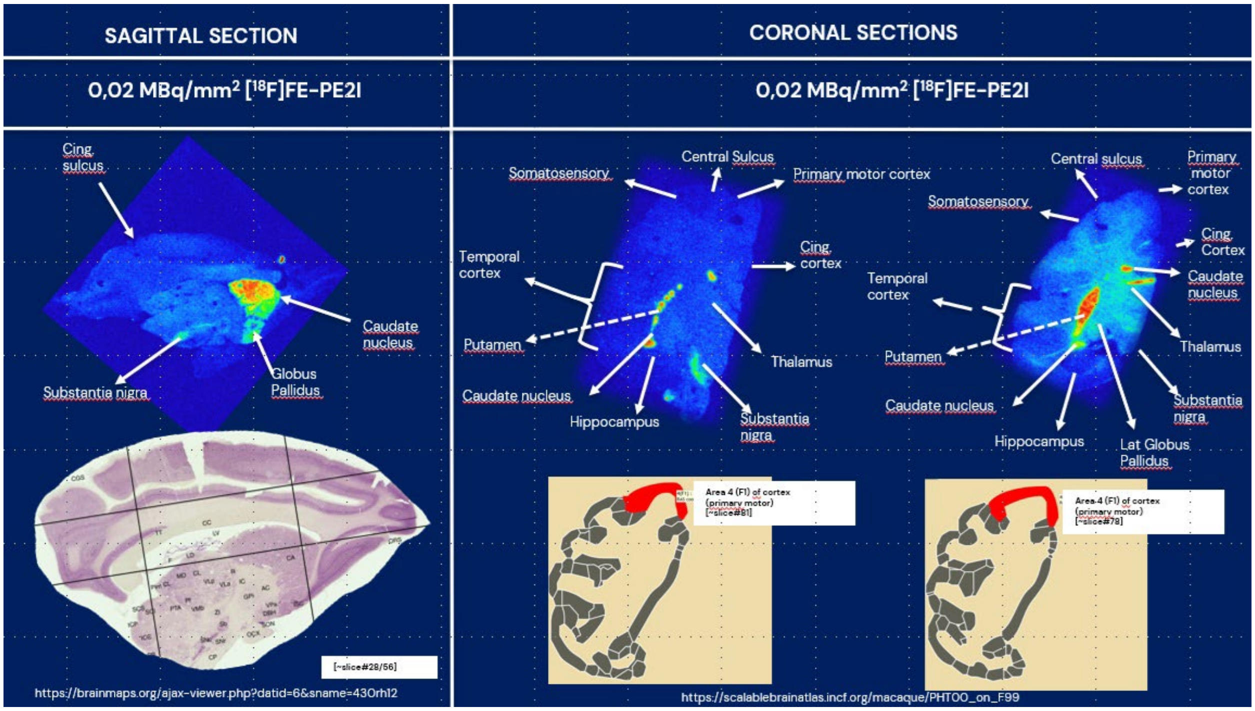


**Figure S6**. Autoradiography study with [$^{18}$F]FE-PE2I on sagittal and coronal sections of three non-human primates.

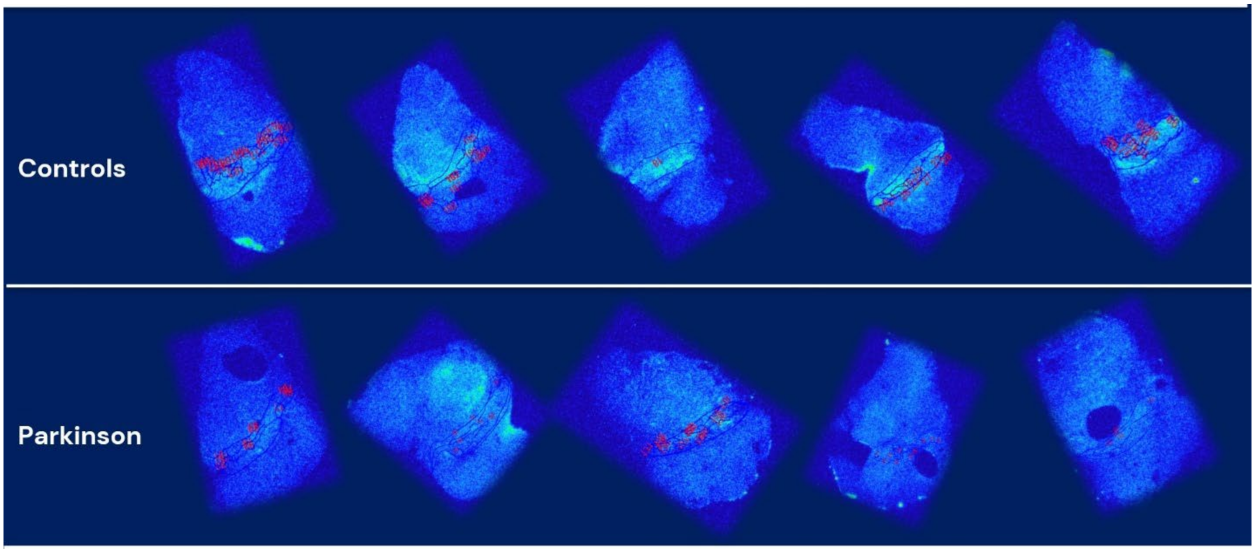


**Figure S7**. Autoradiography study with [$^{18}$F]FE-PE2I on transaxial brain tissue sections from five control donors and five donors with Parkinson´s disease.

**A**

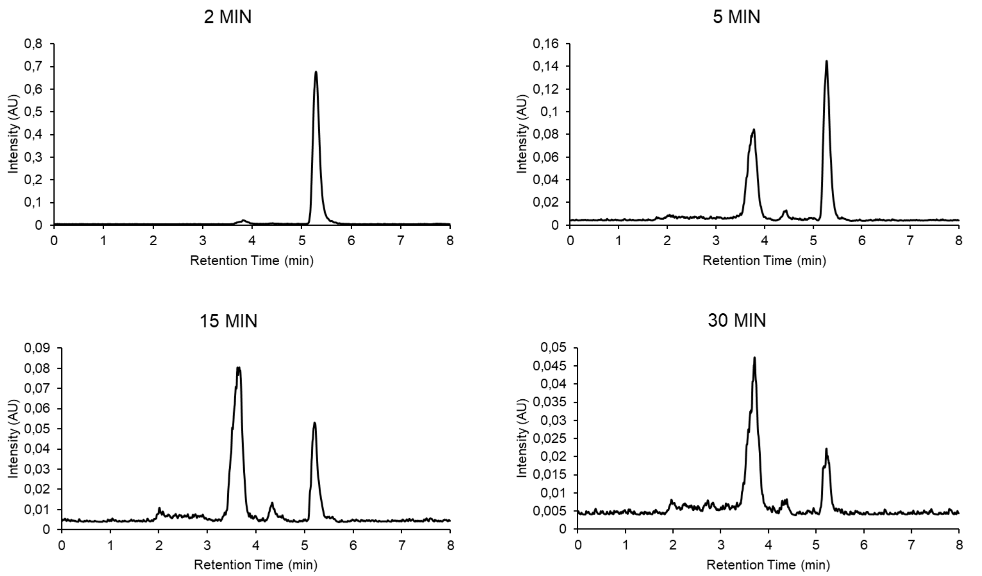


**B**

| | | | % DRM T120 | | | |
|---|---|---|---|---|---|---|
| *RRT* | *M+H* | | Human | Monkey Cyno | Rat | Mouse |
| *0.70* | *351.12405* | M1__+(O2) | | 1% | 6% | 0% |
| *0.73* | *367.11903* | M2_pyran or N,S-ox (O2)_+(O3) | | 2% | | 2% |
| *0.75* | *367.11905* | M3__+(O3) | 1% | 1% | | 2% |
| *0.76* | *337.14458* | M4_pyran or N,S-ox_+(H2 O) | 0% | 2% | | 0% |
| *0.76* | *335.12905* | M5_Et _+(O) | 4% | 5% | 3% | 0% |
| *0.80* | *351.12405* | M6_pyran or N,S-ox_+(O2) | 27% | 76% | 35% | 72% |
| *0.87* | *349.10843* | M7_pyran_-(H2) +(O2) | | 7% | 16% | 4% |
| *0.92* | *335.12915* | M8_pyran or N,S-ox_+(O) | 0% | 6% | 40% | 20% |
| *1.00* | *319.13411* | UCB1436020 | 68% | 0% | 0% | 0% |

**C**

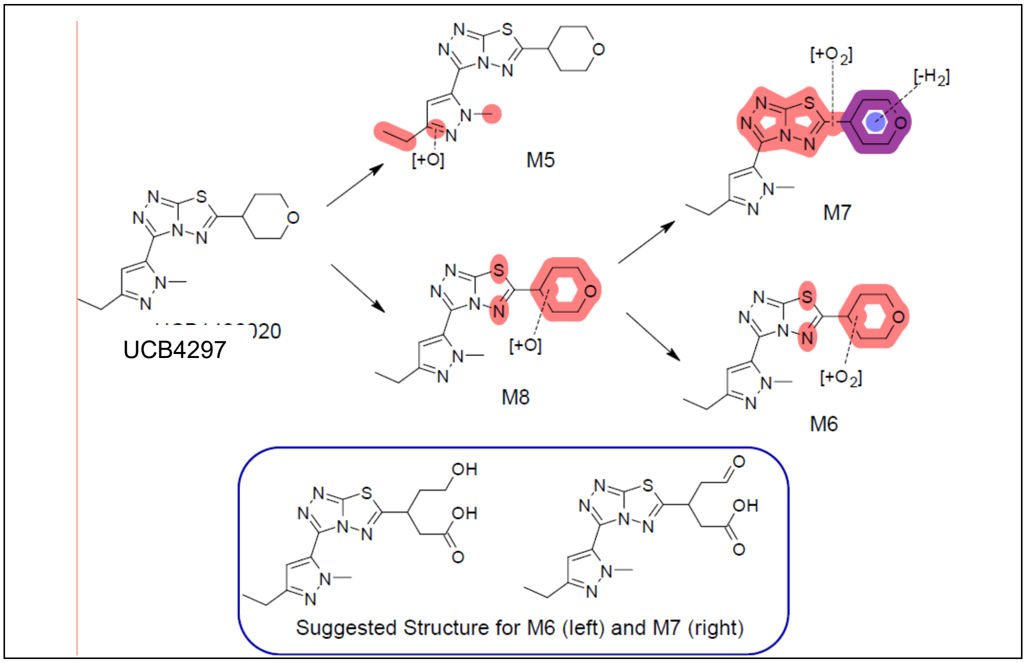


**Figure S8**. A) Representative HPLC chromatogram of the analysis of [$^{11}$C]UCB-1A plasma radioactivity at different time points post-injection; B) Mass spectrometry studies in primary hepatocytes from Caucasian human, Cynomolgus monkey, Sprague Dawley rat and Balb/c mouse. B) Suggested metabolic pathway of UCB-1A (UCB4297).

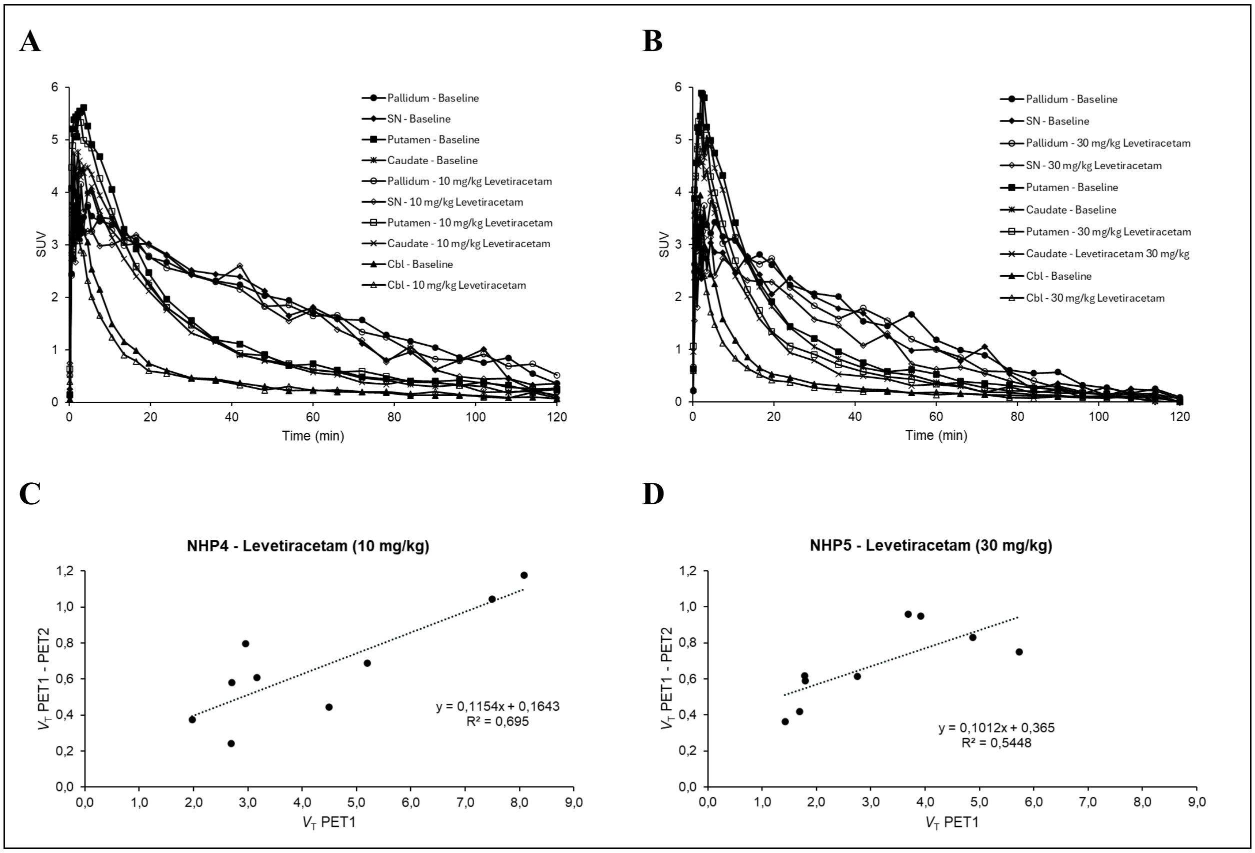


**Figure S9**. Time-activity curves of [$^{11}$C]UCB-1A in caudate, putamen, pallidum, substantia nigra and cerebellum before and after intravenous administration of A) 10 mg/kg Levetiracetam in NHP4 and B) 30 mg/kg Levetiracetam in NHP5. Corresponding revised Lassen plot for the doses of Levetiracetam C) 10 mg/kg in NHP4 and D) 30 mg/kg in NHP5.

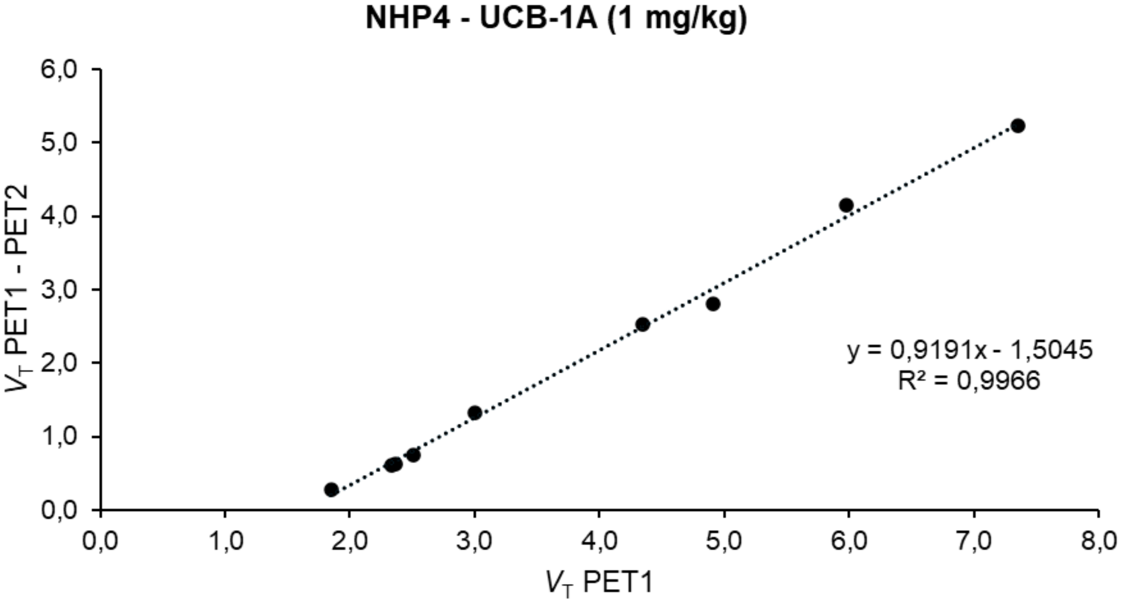


**Figure S10.** Revised Lassen Plot for estimation of occupancy of UCB-1A on the binding of [[11]C]UCB-1A to SV2C.

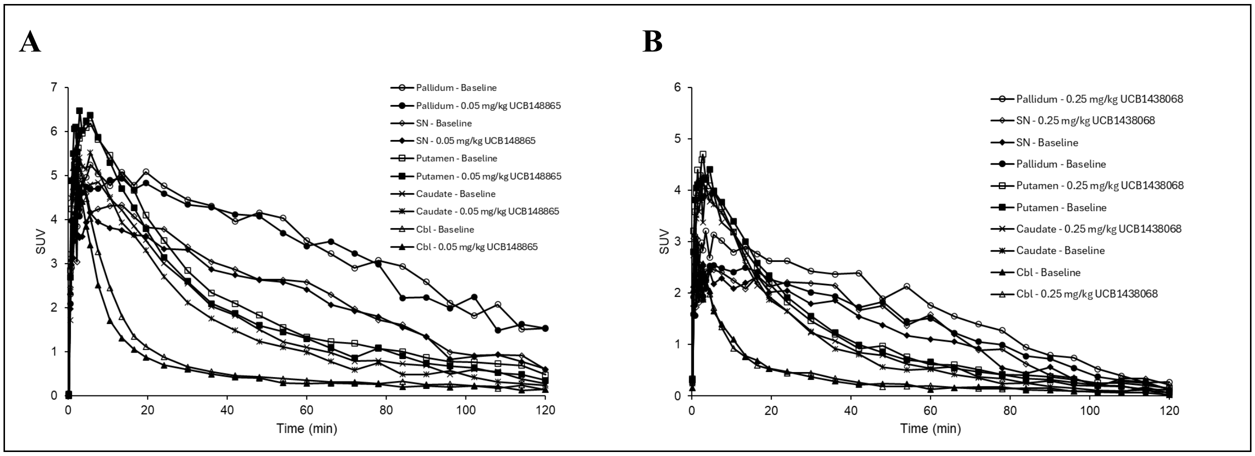


**Figure S11**. Time-activity curves of [[11]C]UCB-1A in caudate, putamen, pallidum, substantia nigra and cerebellum before and after intravenous administration of A) 0.05 mg/kg Seletracetam (SV2A blocker) in NHP6 and B) 0.25 mg/kg UCB5203 (SV2B blocker) in NHP5.

**Table S1**. Structure and affinity values of two selective hits from UCB library.

| Hit nr | Structure | $pIC_{50}$ hSV2C | $pIC_{50}$ hSV2A/B |
|---|---|---|---|
| **1** | 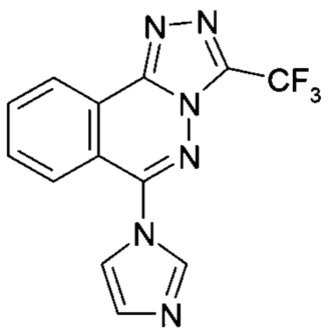 | 7.3 | 5.4/6.6 |
| **1a** | 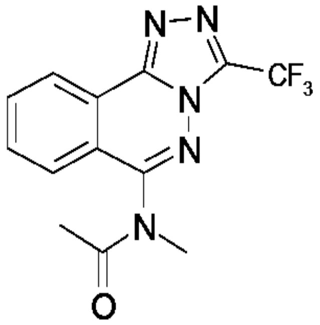 | 6.8 | <5.5 |
| **1b** | 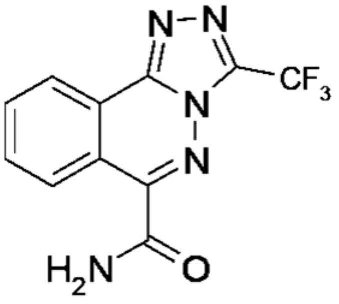 | 7.4 | <5/5.8 |
| **2** | 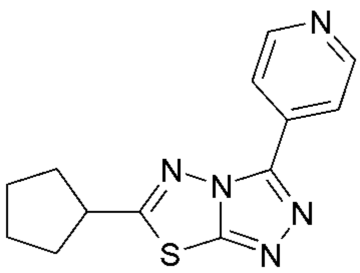 | 7.7 | 6.7/6.1 |

**Table S2**. Weight of non-human primates and experimental details of the PET studies with [$^{11}$C]UCB-1A.

| **NHP#** | **Weight (kg)** | **Injected radioactivity (MBq)** | **Molar activity (GBq/µmol)** | **Injected mass (µg)** | **Condition** | **Blood sampling** |
|---|---|---|---|---|---|---|
| NHP3 | 7.8 | 153 | 263 | 0.18 | Baseline | Venous |
| NHP2 | 9.8 | 161 | 285 | 0.18 | Baseline | Venous |
| NHP4 | 5.7 | 160 | 323 | 0.16 | Baseline | Arterial |
| | | 144 | 433 | 0.11 | Levetiracetam (10 mg/kg) | Arterial |
| NHP5 | 6.8 | 150 | 114 | 0.42 | Baseline | Arterial |
| | | 156 | 192 | 0.26 | Levetiracetam (30 mg/kg) | Arterial |
| NHP5 | 7.4 | 186 | 504 | 0.12 | Baseline | Venous |
| NHP4 | 6.2 | 189 | 1194 | 0.05 | UCB-1A (1 mg/kg) | Arterial |
| NHP5 | 7.4 | 197 | 320 | 0.20 | UCB-1A (1 mg/kg) | Venous |
| NHP4 | 6.3 | 204 | 460 | 0.14 | Baseline | Arterial |
| NHP5 | 6.8 | 190 | 299 | 0.20 | Baseline | Arterial |
| | | 212 | 294 | 0.23 | UCB1438068 (0.25 mg/kg) | Arterial |
| NHP6 | 7.1 | 190 | 331 | 0.23 | Baseline | Arterial |
| | | 187 | 315 | 0.18 | UCB148865 (0.05 mg/kg) | Arterial |

**Table S3**. Binding potential of [$^{11}$C]UCB-1A using $V_{ND}$ estimated with revised Lassen plot.

| Region | Baseline | UCB-1A 1 mg/kg | Occupancy SV2C (%) | Baseline | Levetiracetam 10 mg/kg | Occupancy SV2A (%) |
|---|---|---|---|---|---|---|
| **Caudate** | 1.65 | 0.11 | 93% | 1.55 | 1.47 | 16% |
| **Putamen** | 2.00 | 0.28 | 86% | 2.17 | 1.75 | 19% |
| **Globus pallidus** | 3.49 | 0.30 | 91% | 3.94 | 3.22 | 18% |
| **Substantia nigra** | 2.65 | 0.12 | 96% | 3.58 | 2.94 | 18% |
| **Cerebellum** | 0.13 | -0.04 | 132% | 0.21 | -0.02 | 111% |

**Table S4**. In vitro pKi (n=3) of UCB-1A to SV2 proteins measured at 4°C and 37°C.

| Radioligand | hSV2C (4°C) | hSV2C (37°C) | rSV2C (37°C) | pSV2C (37°C) | hSV2A (37°C) | hSV2B (37°C) |
|---|---|---|---|---|---|---|
| **[$^{3}$H]UCB1275-1** | 8.7 | 8.4 | 8.2 | 8.4 | | |
| **[$^{3}$H]UCB4297** | - | 8.4 | 8 | 8.2 | | |
| **[$^{3}$H]padsevonil** | - | 8.2 | | | <6 | 6.1 |

[$^{3}$H]UCB1275-1: from Shi J, et al. Biochemical Society Trans 2011;39,5:1341-7.
[$^{3}$H]UCB4297=[$^{3}$H]UCB-1A

**Movie S1**

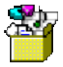

11C-UCB-1A NHP PET.mp4
